# Reversible Dissociation of Mitochondrial Complex V Balances Anabolic and Energy-Generating Needs in Cancer

**DOI:** 10.1101/2025.08.05.668642

**Authors:** Huabo Wang, Jie Lu, Colin Henchy, Steven J. Mullett, Adam C. Richert, Eric S. Goetzman, Ahmet Toksoz, Leonardo Hale, Brandon B. Wang, Nelli Mnatsakanyan, Edward V. Prochownik

## Abstract

Cancer cell metabolic re-programming provides the excess energy and anabolic precursors necessary to sustain uncontrolled growth. This is partly mediated by the Warburg effect, whereby glucose is converted into ATP and a subset of these anabolic substrates. Concurrently, mitochondrial mass and ATP production decline in most tumors. This raises the question of how increased supplies of glycolysis-derived anabolic substrates can be balanced with those generated by the TCA cycle. Using primary murine liver cancers and cell lines, we show that this can be explained by the dissociation of mitochondrial Complex V (CV or ATP synthase) into its component and functionally-independent F_o_ and F_1_ domains. This occurs as a result of marked reductions in MT-ATP6, a CV subunit that stabilizes the F_o_-F_1_ association. Serving as a proton pore, F_o_ maintains a normal mitochondrial membrane potential without generating ATP, thus allowing the TCA cycle, electron transport chain and anaplerotic reactions to function at high levels. Concurrently, free F_1_ functions as an ATPase to prevent excessive ATP accumulation. The uncoupling of TCA cycle-derived anabolic substrate production from membrane hyperpolarization and ATP synthesis by a smaller population of more efficient mitochondria allows TCA cycle-generated anabolic precursors to match those generated via glycolysis.

## Introduction

Cancer cells are confronted with numerous metabolic challenges that require rapid adjustments in the sources from which they generate ATP and how it is allocated. To a large degree, this reflects the energy-demanding processes of enhanced translation and biomass accumulation associated with neoplastic proliferation ^1–3^. Certain oncoproteins such as MYC contribute to this metabolic re-programming by increasing the expression of genes that regulate ATP production via glycolysis and oxidative phosphorylation (Oxphos), the anabolic substrates derived from these two sources and the components and activities of the translational machinery ^4–11^.

Although the benefits of tumor-specific metabolic re-adjustment are well-known, a failure to coordinate glycolysis and Oxphos can also negatively impact growth if it produces imbalances in ATP production and/or anabolic substrate supplies. For example, excessive ATP is toxic by virtue of chelating magnesium and limiting the cation’s roles in maintaining ribosomal structural and functional integrity ^12^. Indeed, the need to constrain ATP levels within a narrow range is believed to be a major reason for the apparent inverse relationship that exists between the activities of the glycolytic pathway and TCA cycle in most cancers ^13–16^. However, this apparent mutual exclusivity raises questions that cannot be satisfactorily answered by the current paradigm of tumor metabolism. Among these is how an optimal balance of anabolic precursors can be maintained in the face of the high levels of glycolysis-derived substrates provided by Warburg-type respiration and the seemingly reduced supplies provided by the TCA cycle. Another question is why, in their presumably diminished role as a source of ATP, tumor mitochondria nonetheless often expand their choice of TCA cycle precursors to include more energy-rich substrates such as fatty acids and increase their reliance on anaplerotic substrates such as glutamine ^2,17–19^. Indeed, anaplerosis itself, which is defined as a replenishing of depleted TCA cycle intermediates would seem to be incompatible with not only the reduced mitochondrial mass of most tumors but with the above-mentioned barriers that restrict their generation of ATP ^14,20,21^.

Mitochondrial ATP production relies upon an electro-chemical proton gradient that generates a membrane potential (ΔΨm) between the mitochondrial inter-membrane space (IMS) and the matrix that is maintained by the electron transport chain (ETC). The electrons powering the ETC are provided in the form of NADH and FADH2 that are generated from TCA cycle substrate oxidation in the matrix during the reduction of the electron acceptors NAD^+^ and FAD^+^. High matrix levels of TCA cycle substrates, protons, NADH and ATP are all potent inhibitors of Oxphos and thus serve as safeguards against hyperpolarization ^22–24^. In turn, low rates of Oxphos reduce the supplies of critical TCA cycle-derived anabolic substrates such as aspartate and acetyl coenzyme A (AcCoA) that are necessary for the *de novo* synthesis of purine and pyrimidine nucleotides, proteins and lipids ^23,25^.

The mitochondrial ATP synthase, also known as Complex V (CV), consists of 2 physically connected domains, F_o_ and F_1_, that coordinate proton re-entry into the matrix from the IMS via the inner mitochondrial membrane (IMM)-embedded F_o_ while driving ATP generation via matrix-localized F_1_ ^26–29^. F_o_ and F_1_ are connected by 2 multi-subunit “stalks”. The central stalk allows the c-ring of F_o_ to rotate in a proton-driven counterclockwise manner relative to F_1_, thus providing the electro-mechanical energy for ATP generation by the latter domain. The peripheral stalk helps to anchor F_o_ to the IMM and maintain its association with F_1_ via the stabilizing influence of the MT-ATP6 (also known as subunit a) and MT-ATP8 subunits, with the former also contributing to F_o_’s function as the proton channel ^27,29–32^. MT-ATP6 and MT-ATP8 are co-translated from a single transcript in partially overlapping reading frames and are the only CV subunits encoded by the mitochondrial genome ^33–36^.

Inherited missense, frameshift or nonsense mutations in MT-ATP6, and less commonly MT-ATP8, occur in a subset of individuals with Leigh syndrome; these destabilize the F_o_-F_1_ connection and render CV susceptible to dissociation ^33,36–39^. Depending upon the nature of the mutation, the level of heteroplasmy and the tissue examined, this causes variable amounts of proton leak through free F_o_, ΔΨm dissipation and reduced ATP synthesis ^40^. Clinical manifestations include variable degrees of ataxia, neuropathy and myopathy often in association with lactic acidosis as cells attempt to compensate for the energy deficit by increasing glycolysis, although not to an extent that fully normalizes ATP levels ^36,37,39^.

Here we describe how, under certain circumstances, tumor cells reversibly down-regulate MT-ATP6 levels, thus creating an imbalance among CV subunits and destabilizing CV in a way that allows for partial F_o_-F_1_ dissociation in a manner that mimics Leigh syndrome. The resulting free F_o_ thus provides a channel through which protons can return to the matrix without generating ATP while free F_1_ can function in reverse as an ATPase to limit ATP over-accumulation and TCA cycle inhibition. In concert, this allows the TCA cycle and ETC to operate more rapidly but without being inhibited by membrane hyperpolarization or an excess of ATP. This allows TCA cycle-derived anabolic substrates to be produced at levels that are now better matched with those derived via the high rates of Warburg effect-related glycolysis ^13,15^. Increased TCA cycle efficiency thus allows for the reduction in mitochondrial mass seen in most tumor cells as well as the marked anaplerosis and altered energy generating substrate preferences that often distinguishes them from their untransformed counterparts.

## Results

### CV partially dissociates into its component F_o_ and F_1_ domains in murine liver cancer models

Non-denaturing gel electrophoresis (NDGE) and *in situ* ATPase assays were used to interrogate and compare the CV activities in normal murine livers and hepatoblastomas (HBs) generated by enforcing the expression of the oncogenic patient-derived β-catenin mutation Δ90 (B) and the yes-associated protein (YAP) mutant S127A (Y) ^41–44^. Together, these generated “BY” HBs that histologically and molecularly closely resemble the crowded fetal subtype of human HB ^42–44^. Both control livers and primary BY HBs showed strong bands of high molecular weight ATPase activity corresponding to the location of intact CV in Coomassie Blue-stained companion gels, whereas most HBs showed additional activity in more rapidly migrating bands (“Free ^7^ATPase”) (Figure 1A and data not shown) ^43,45,46^. This was also seen in BY HBs generated in livers bearing hepatocyte-specific inactivation of the genes encoding pyruvate dehydrogenase (*Pdha1*) and the carbohydrate response element binding protein (*Chrebp*), a Myc-related transcription factor that is a member of the extended Myc network (Supplementary Figure 1) ^10,41,47^. Similar findings were made in a mouse model of highly undifferentiated hepatocellular carcinoma (HCC) driven by the doxycycline-regulatable over-expression of a human *MYC* transgene (Figure 1B) ^45^. Although the free ATPase activity was much less prominent in MYC-over-expressing livers that had yet to develop tumors, it persisted for at least 3 days following *MYC* deactivation and the onset of tumor regression and then re-appeared in recurrent tumors (Figure 1B).

**Figure 1.**
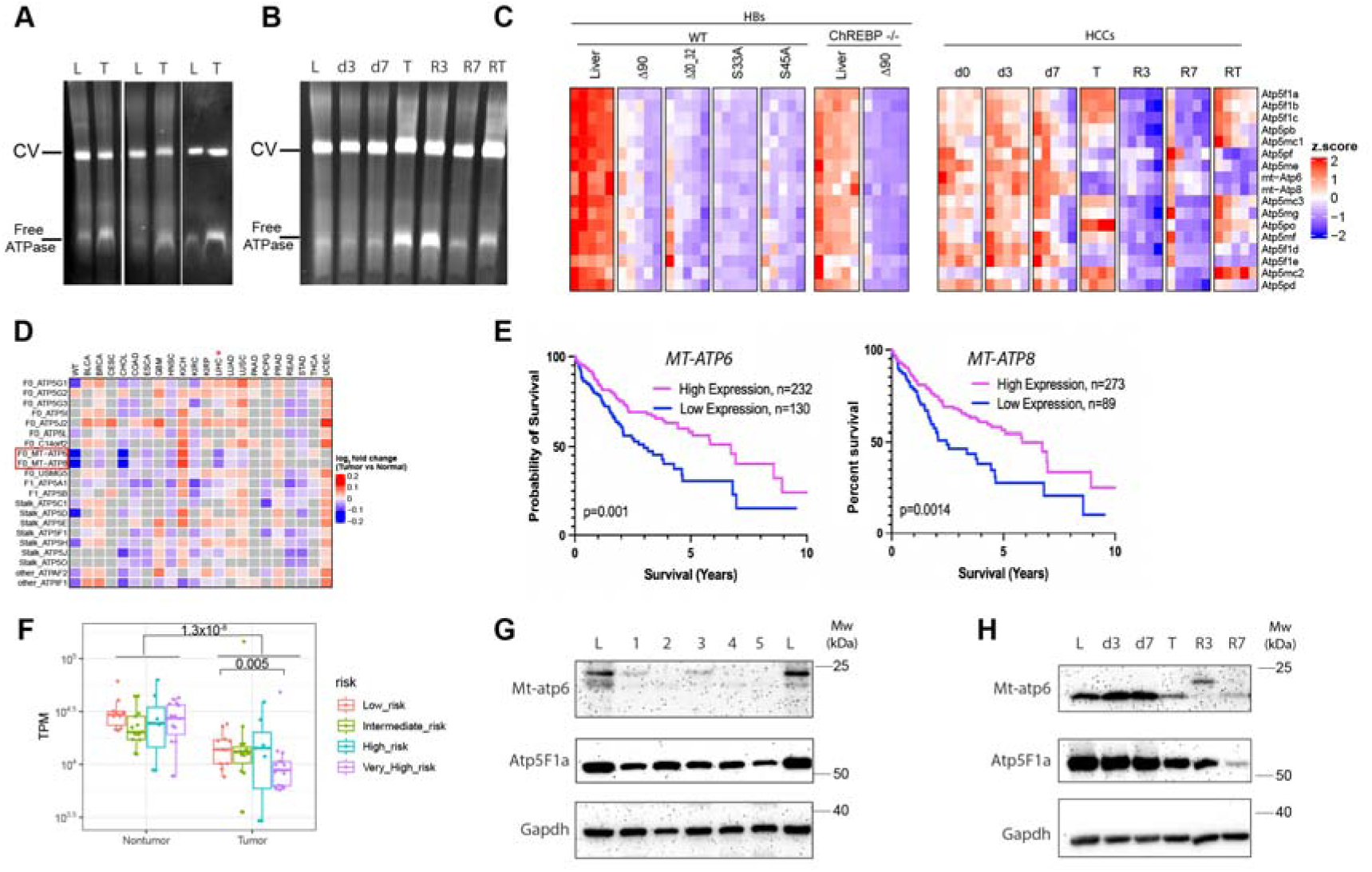
HBs and HCCs express “free ATPase” and down-regulate transcripts encoding *MT-ATP6, MT-ATP8* and other CV subunits. (A). NDGE and *in situ* ATPase assays performed on ETC complexes from livers (L) and HBs (T) ^7,43,44^. In all cases, enzyme activity was detected in the region corresponding to the known location of CV ^7,43–45^. Tumors also contained significantly larger amounts of an additional, more rapidly migrating ATPase activity (“Free ATPase”). (B). NDGE and *in situ* ATPase assays performed on ETC complexes from murine livers and HCCs generated by a doxycycline-regulated human *MYC* transgene. Liver samples on days 0, 3 and 7 after MYC induction (D0, D3 and D7) were obtained prior to the appearance of visible HCCs (lanes 1-3) ^45^. On day ∼30, large HCCs (T) were now present at which point *MYC* expression was silenced and regressing tumors were sampled 3 and 7 days later (R3, R7). Finally, ∼3 months after tumors had completely regressed, *MYC* was again induced and a recurrent tumor (RT) was sampled ∼30 days later. (C). Expression of CV subunit-encoding transcripts in murine livers and BY HBs and *MYC*-induced HCCs ^43–45^. The former included tumors generated with several patient-derived β-catenin mutants as well as tumors induced by B in *Chrebp-/-* hepatocytes). Note some strain- and analysis-specific variation among the livers from HB and HCC groups. Each column represents transcripts from one of 4-5 individuals in the indicated group. Blue bracket: transcripts that are selectively down-regulated in HBs and in primary and recurrent HCCs and that encode members of the central and peripheral stalks. Red bracket: *Mt-atp6* and *Mt-atp8* transcripts. (D). Expression of CV subunit-encoding transcripts among human cancers in TCGA. Red star: HCCs (LIHC); Red box: *MT-ATP6* and *MT-ATP8* transcripts. (E). Kaplan-Meier survival of HCC/LIHC patients from (D) based on *MT-ATP6* and *MT-ATP8* transcript levels. (F). Low levels of *MT-ATP6* and *MT-ATP8* transcripts are associated with high-risk human HBs (data from Carillo-Reixach and Hooks ^55,56^). (G). Immunoblot for Mt-atp6 protein in control mouse livers (L) and 5 representative HBs (1-5) similar to those in A. (H). Immunoblot for Mt-atp6 protein during the course of murine HCC induction and regression as depicted in (B).

### Murine liver cancers and many human tumors down-regulate *MT-ATP6* and *MT-ATP8* transcripts and cause CV subunit imbalance

To investigate the basis of the above-noted free ATPase activity, we surveyed RNAseq data from additional murine HBs and HCCs, focusing initially on transcripts that encode CV subunits ^26,28^. In the former case, we included data from HBs generated with several patient-derived β-catenin mutants other than B (Δ90) as well as those generated in the above-mentioned *Chrebp-/-* mice ^43^. We paid particular attention to transcripts encoding central and peripheral stalk proteins and MT-ATP6 and MT-ATP8, which stabilize the F_o_-F_1_ interaction ^27,29,33,48,49^ (Figure 1C). All HBs down-regulated *MT-ATP6* and *MT-ATP8* transcripts as well as all other CV-related ones whereas primary and recurrent HCCs showed more selective and complex changes (Figure 1C). For example, *Atp5f1a* and *Atp5f1b* transcripts, which encode the α and β major structural subunits of F_1_, with β containing the catalytic nucleotide binding sites, were actually up-regulated in both primary and recurrent tumors. *Atp5po,* which encodes OSCP, the major component of the peripheral stalk, was also upregulated. In contrast, *Atp5f1d* and *Atp5f1e* transcripts, which respectively encode the δ and ε central stalk subunits, as well as *Mt-atp6* and *Mt-atp8* transcripts, were down-regulated. These findings suggested that free ATPase activity (Figure 1A and B) might be a consequence of upregulation of F_1_ subunit levels (since the minimal unit required for ATP hydrolysis is the α3β3γ subcomplex), as well stoichiometric imbalances among CV’s central and/or peripheral stalk subunits and stabilize the F_o_-F_1_ association.

We then focused more specifically on Mt-Atp6 and Mt-Atp8 for 3 reasons. First, as the only CV subunits encoded by the mitochondrial genome and specifically down-regulated in HCCs, they are co-translated from a single mature transcript in partially overlapping reading frames ^33,50–52^. Accordingly, their expression is similarly regulated. Second, models of CV assembly have shown that the Mt-Atp6 and Mt-Atp8 subunits are added to and stabilize CV after F_o_ and F_1_ are already connected via the central and peripheral stalks^26–29,49^. Third, inherited missense mutations, primarily involving *MT-ATP6*, occur in a subset of individuals with Leigh syndrome in association with variable degrees of F_o_-F_1_ dissociation, lactic acidosis and multiple clinical manifestations arising as a result of inadequate ATP production ^34,50,53,54^.

*MT-ATP6* and *MT-ATP8* transcripts were among the most consistently and coordinately down-regulated and/or imbalanced of all CV-related mRNAs across human HBs and HCCs and other evaluable human tumor types in the TCGA database (Figure 1D and Supplementary Figures 2 and 3) ^55,56^. HCCs expressing the lowest levels of these transcripts were also associated with significantly shorter survival whereas in HBs, low-level expression was associated with “very high risk” features (Figure 1E and F and Supplementary 2). Finally, comparative immunoblot analyses of murine livers, BY HBs and Tet-MYC-induced HCCs showed a marked down-regulation of Mt-atp6 protein in both tumor types (Figure 1G and H).

### Murine HBs and HCCs reduce their mitochondrial mass but increase TCA cycle activity

We next asked whether the HB- and HCC-associated free ATPase activity identified in Figures 1A and B and Supplementary Figure 1 might originate from CV’s F_1_ domain that had dissociated from F_o_ as a result of reduced amounts of Mt-atp6 and CV destabilization. Functionally, quantitative changes in Mt-atp6 seen in Figure 1G and H would be predicted to mimic the qualitative and functionally inactive ones associated with Leigh syndrome ^36,37,39,53^. This might allow for increased TCA cycle and ETC activity in tumors by permitting free F_o_ to assume a secondary role as a mitochondrial permeability transition pore (mPTP). Transient openings of mPTP that serve this function are known to prevent hyperpolarization^31^, thereby maintaining a normal ΔΨm in a manner that neither generates ATP nor inhibits the ETC ^31,32,57,58^. Finally, and if necessary, free F_1_ could function as an ATPase that would limit the toxicities associated with any excessive ATP that was produced while maintaining levels sufficient to meet proliferative demands ^1,12,59^.

To test these ideas, we first compared the mitochondrial mass and TCA cycle function of normal livers and BY HBs plus those generated by the pairwise combination of Y^S127A^ and 13 other patient-derived β-catenin mutants ^43^. Using 2 different TaqMan probes to quantify mitochondrial DNA (mtDNA) we confirmed previous findings that HBs, like most other human cancers, have a ∼6-fold lower mitochondrial mass relative to matched normal tissues (Supplementary Figure 4A) ^14,43,45^. In contrast, Complex I (CI) activities of all HBs, as measured by oxygen consumption rates (OCRs) in response to pyruvate, malate, glutamate and ADP, were significantly higher even before adjusting for differences in mtDNA mass differences (Supplementary Figure 4B). These HBs displayed even higher relative rates of pyruvate oxidation to acetyl-CoA (Supplementary Figure 4C), which was consistent with our previous observation that they also contained significantly higher levels of active pyruvate dehydrogenase complex activity and AcCoA content ^44^. We have previously reported similar findings with regard to mtDNA content and PDH activity in MYC-induced HCCs ^45^.

The finding that HBs and HCCs have reduced levels of CV subunit-encoding transcripts despite maintaining high levels of TCA cycle activity (Figure 1) ^43–45^ was extended to transcripts encoding the majority of subunits of the mitochondrial respiratory chain (i.e. Complexes I-IV and the remaining mitochondrial proteins (Supplementary Figure 4D-G). Overall, these findings are consistent with those pointing to significant reductions in mitochondrial mass and/or widespread imbalances among mitochondrial-specific transcripts and proteins despite up-regulation of TCA cycle activity ^60^.

### Free ATPase corresponds to CV’s F_1_ domain and its association with F_o_ can be regulated

To unequivocally establish the identity of free ATPase and its metabolic and biological ramifications in greater detail and under more controllable circumstances, we derived immortalized cell lines from BY HBs ^61^. Two such lines, BY1 and BY3, were identified that either did or did not contain the same free ATPase activity identified in the primary tumors from which they were derived (Figure 2A). Like the original tumors, both cell lines maintained low levels of Mt-atp6 expression (Figure 2B). Possibly explaining why BY3 cells did not contain free F_1_ was their lower levels of ATP5F1A (i.e. the α subunit of F_1_), which might have re-balanced CV’s aberrant subunit stoichiometry and allowed it to better maintain the F_o_-F_1_ association. A longer period of *in situ* ATPase assay development revealed additional bands of free ATPase activity associated with BY1 cells (Figure 2C). Mass spectrometry analysis revealed that the slowest migrating (top) band contained all CV proteins, that the one just beneath it contained the δ, γ and ε components of the central stalk plus the α and β subunits of F_1_ and that the fastest migrating band contained only the α and β subunits that comprise the minimally active F_1_ domain ^26,62^. In neither of these latter 2 bands did we detect ATP5IF1/IF1, which is a potent inhibitor of both the ATP synthase and ATPase activity of free CV and free F_1_, respectively that also ^63,64^.

**Figure 2.**
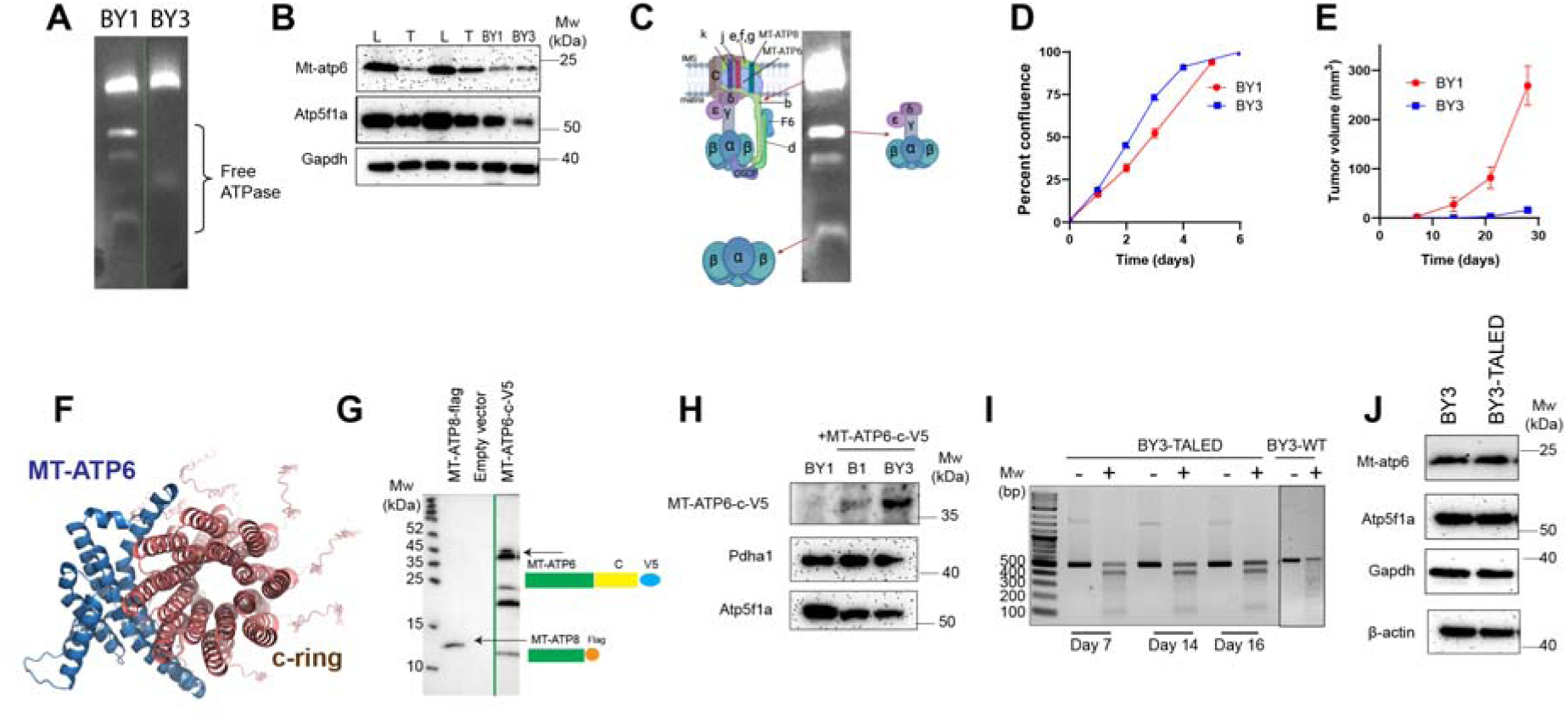
Free ATPase is comprised of the F_1_ domain and can be genetically manipulated. (A). NDGE and *in situ* ATPase assays performed on ETC complexes isolated from BY1 and BY3 HB cell lines ^61^. (B). Immunoblot analysis for Mt-atp6 and Atp5f1a (α subunit of CV) in BY1 and BY3 cells. Control liver and BY HB lysates were used as controls to show that the cell lines retained the low-level expression of Mt-atp6 associated with tumors (Figure 1G). (C). Over-exposure of the ATPase assay performed in (A). The adjacent cartoons depict the CV-related proteins that were identified in the bands excised from the indicated regions of the gel using qualitative protein MS. (D). *In vitro* growth curves of BY1 and BY 3 cells. Each point represents the mean of 6 replicas +/- 1 S.E. (E). BY1 and BY3 tumor growth rates. 10^6^ cells of each type were grown as subQ tumors in FVB mice. Tumor volumes were measured at the times indicated. n=4 mice/group. (F). Structural relationships between Mt-atp6 and the c-subunit ring (from ref. 27). The cradling of the c-ring by the α5 and α6 helices of Mt-atp6 forms part of the proton channel ^29,94^. (G). Mt-atp8 and Mt-atp6 immunoblots in BY1 cells. Lane 2: total lysate from BY1 cells stably expressing murine Mt-atp8-Flag tag protein. The precursor protein (not seen) contained a mitochondrial targeting sequence (MTS) at its N-terminus. Only the completely processed C-terminal Flag-tagged protein is seen. Lane 3: total lysate of BY1 cells stably transfected with a control, empty SB vector. Lane 4: Total lysate of BY1 cells transiently transfected with a SB vector encoding a murine Mt-atp6-c subunit-V5 fusion protein. As with the Mt-atp8 protein, no precursor protein containing the localization signal was detected. Different regions of the blot were probed with anti-FLAG and anti-V5 antibodies to allow for detection of both MT-ATP8-FLAG and MT-ATP6-V5 proteins. (H). Stable expression of Mt-atp6-c-V5 fusion protein from A in purified BY1 and BY3 mitochondria. Lane 1: a mitochondrial lysate from WT BY1 cells transfected with an empty vector served as a negative transfection control. The Mt-atp6-c blot was probed with an anti-V5 antibody. (I). TALED-generated mutations of the *Mt-atp6* gene. BY3 cells were transiently transfected with TALED vectors and a control vector expressing GFP. The EGFP+ population was then purified by FACS on day 2 and evaluated for *Mt-atp6* heteroplasmy at the times indicated. For this, a 504 bp fragment of mtDNA spanning the *Mt-atp6* gene coding sequence was amplified from BY3-TALED cells or WT-BY3 cells. The fragments were melted, re-annealed and either digested with T7 endonuclease (+) or not (-) followed by 2% agarose gel electrophoresis. Sequencing of the PCR products obtained on ∼day 23 documented the expected mutations at a total frequency of 37% (Supplementary File 1). (J). Immunoblots of endogenous Mt-atp6 protein in WT BY3 and BY3-TALED cells on day 14.

BY1 and BY3 cells showed minimal differences in their *in vitro* proliferation rates but large differences when propagated as subQ tumors, with BY1 tumors having a substantial growth advantage (Figure 2D and E). To examine the relationship between growth rate and free F_1_ more closely, we first attempted to enforce the ectopic expression of Mt-atp6 in BY1 cells and thus restore F_o_-F_1_ re-association. However, the extreme hydrophobicity of Mt-atp6 made its expression challenging to document as previously reported by several other groups (not shown) ^33,51,52,65,66^. We therefore explored alternative approaches to overcome this barrier and ultimately capitalized on Mt-atp6’s intimate relationship with F_o_‘s c subunit by expressing a Mt-atp6-c-V5 epitope-tagged fusion protein (hereafter dubbed Mt-atp6-c) (Figure 2F and G) ^29^. Mt-atp6-c was both readily detectable and localized to mitochondria in both BY1 and BY3 cells (Figure 2H). *In situ* ATPase assays performed on these “BY1-Mt-atp6-c” cells showed a significant reduction in free F_1_ subunit levels and no effect in BY3 cells (Supplementary Figure 5A and not shown). Along with the normalization of ATPase activity that is a property of F_1_’s β subunit, BY1-Mt-atp6-c cells also showed a large reduction in the amount of co-migrating α subunit (Supplementary Figure 5B). Thus stably expressing Mt-atp6-c rescued most of the CV defect of BY1 cells.

In a complementary approach to manipulating free F_1_ levels we used mitochondrial-targeted TALE-linked deaminases (TALEDs) ^67^ to allow for site-specific A→G base editing of mtDNA and the generation of 3 *Mt-atp6* missense mutations two of which (L220P and L222P) have been previously described in Leigh syndrome and one of which (Y221H) has not ^39,68^. These remained stable throughout the study and, on average, comprised ∼37% of all *Mt-atp6* sequences, which was in excellent agreement with the previously reported level of stable heteroplasmy achieved in murine embryos (Figure 2I and Supplementary Table 1) ^67^. These mutations did not affect Mt-atp6 protein levels but did increase the amounts of free F_1_ and α subunit as expected (Figure 2J and Supplementary Figure 5C and D).

### F_o_-F1 dissociation provides a proliferative advantage and correlates with increased rates of glycolysis

In both WT BY1 and BY1-Mt-atp6-c cells, we found that hypoxia had little effect on free F_1_ levels (<15% change, not shown). This likely reflected a combination of non-mutually exclusive factors including the differential regulation, abundance and nuclear origin of the Mt-atp6-c fusion protein relative to that of endogenous, mitochondrial-encoded Mt-atp6. In contrast, WT BY3 cells demonstrated significant increases when maintained for 2 days under hypoxic conditions and BY3-TALED cells increased this level by an additional 2-3-fold (Figure 3A). Hypoxia also stimulated the uptake of the glucose analog NBDG and supported the idea that increased supplies of glycolysis-derived anabolic substrates were being balanced by reciprocal increases in TCA cycle-derived substrates as a result of F_o_-F1 dissociation (Figure 3B). Finally, free F_1_ levels impacted the *in vitro* proliferation of BY3 cells in a hypoxic environment. Whereas WT BY3 cells were unable to sustain proliferation beyond 3 days, BY3-TALED cells not only continued to proliferate beyond this point but grew as well as or even better than they did under normoxic conditions and achieved similar densities (Figure 3C).

**Figure 3.**
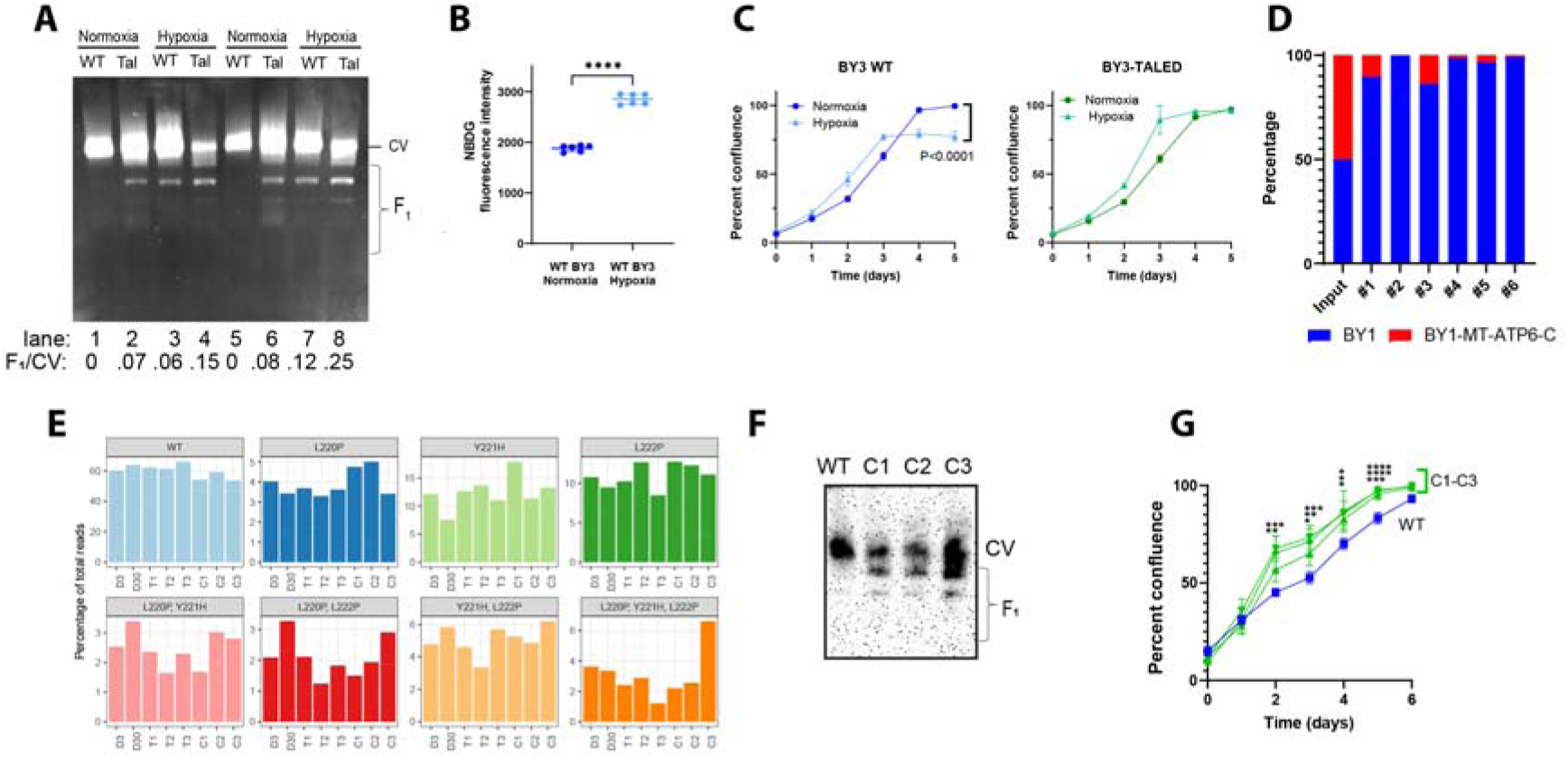
Altering free F_1_ levels impacts free F_1_ levels and proliferation. (A). Duplicate NDGE and ATPase assays performed on BY3 control and BY3-TALED cells. (B). NBDG uptake by BY3 cells under normoxic and hypoxic condition. (C). Growth rates of WT BY3 and BY3-TALED cells under normoxic or hypoxic conditions. Cell density was monitored at the indicated times using an Incucyte SX5 imaging system. Each point represents the mean of 4-6 replicas +/- 1 S.E. (D). Free F_1_ is associated with a proliferative advantage *in vivo.* Equal numbers of WT BY1 + BY1-M-ATP6^cyto-roGFP^ and WT BY1^cto-roGFP^ + BY1-Mtatp6-c (10^6^ of each) were injected subcutaneously into nu/nu mice and grown as tumors for ∼4-5 wks. Total tumor DNAs, along with DNAs from each of the initial 2 mixed populations of input cells were then used in a TaqMan assay to amplify both GFP and human β-catenin. The relative fraction of each population was then determined. (E). Changes in *Mt-atp6* mutation frequencies in BY 3-TALED cells maintained under different conditions. BY3 cells were transfected with TALED vectors that introduced each of the indicated mutations in the *Mt-atp6* gene. PCR amplification of a segment of *Mt-atp6* flanking the mutations was performed on *in vitro*-maintained cells day 3 and day 30 after transfection, on subQ tumors (T1-T3) and on cell lines re-derived from each of the tumors and expanded *in vitro* for an additional 3 wks (Figure 2I). NextGen sequencing was used to establish the frequencies of WT and mutant *Mt-atp6* sequences and the identities of the latter. (F). NDGE of the indicated cell lines followed by transfer to PVDF membrane and immunoblotting for α-subunit. Numbers below the image indicate the fractional amount of non CV-associated α-subunit. (G). Growth curves of WT BY3 cells and BY3-TALED C1-C3 cell lines from (E).

The proliferative advantage of cells with free F_1_ could also be demonstrated *in vivo.* For these studies, WT BY1 and BY1-Mt-atp6-c cells were each stably transfected with a vector that encoded cyto-roGFP as a way of distinguishing the 2 populations. 10^6^ of these cells were then combined with an equal number of untagged cells from the other cell line (i.e. WT BY1 + BY1-M-ATP6^cyto-roGFP^ and WT BY1^cto-roGFP^ + BY1-Mt-atp6-c) and propagated as subQ tumors in nu/nu mice for 5-6 wks. This approach served to control for the possibility that cyto-roGFO non-specifically inhibits growth or elicits a residual immune response that would interfere with the growth of the tagged cells. Regardless of whether they expressed cyto-roGFP or not, BY1 cells always displayed a significant growth advantage over BY1-Mt-atp6-c cells (Figure 3D).

Mitochondrial heteroplasmy is dynamic and subject to tissue-specific changes by factors such as selective DNA replication, aging and cancer ^69–73^. We therefore asked whether time and/or different growth and selection conditions altered the *Mt-atp6* mutational landscape that was initially generated by TALED-mediated mutagenesis. We examined this in BY3 cells maintained *in vitro* for 3 days or 30 days after transiently transfecting the TALED vectors, in 3 subQ tumors (T1, T2, T3) generated by the latter cells and in cell lines that were re-established from each tumor or control tumors originating from WT BY3 cells (C1, C2, C3 and WT). For each sample, the TALED-targeted region of the *Mt-atp6* gene was amplified and subjected to NexGen sequencing as described above to determine the overall frequency of each mutation and its permutations. In addition to displaying mutational frequency of ∼35-46% among the 4 groups, this approach identified samples in which one or more of the mutations was selectively gained or lost. For example, culturing the initially transfected cells for 30 days *in vitro* led to significant increases in the L220P/Y221H, L220P/L222P and Y221H/L222P pairs of mutations indicating that these combinations conferred a growth advantage relative to others (Figure 3E). Conversely, loss of the single L220P and L222P mutations relative to those observed on day 3 suggested that they conferred a selective growth disadvantage. Tumors derived from these cells showed additional selection relative to the day 30 input population, most notably increases in Y221H and decreases in L220P/Y221H and L220P/L222P. Finally, *in vitro* re-growth of the C1-C3 cell lines from these tumors selected for further increases in Y221H and L222P while concurrently selecting for decreases in L220P/Y221H and L220P/L222P. BY3-TALED cells thus undergo non-random albeit complex *in vitro* and *in vivo Mt-atp6* changes in heteroplasmy frequency and content that likely represents selection for the mutational landscapes and the ensuing metabolic adaptations that impart the strongest proliferative advantages. The differences among the mutations from *in vitro-* and *in vivo-*selected cells also indicated that these adaptations are not identical and are selected to maximize growth in different environments.

The C1-C3 cell lines contained significantly larger amounts of free α subunit and grew significantly faster than WT BY3 cells (Figure 3F and G). Thus, cell line-specific adjustments in heteroplasmy content were associated with qualitative and quantitative alterations in free F_1_ and improved proliferative fitness.

### Cells with free F_o_ and F_1_ alter their metabolism while maintaining normal ATP levels and ΔΨm

The foregoing findings indicated that changes in Mt-atp6 levels drive CV’s partial and reversible dissociation into its independently functioning F_o_ and F_1_ domains (Figures 1B and 3B)^74^. Among the properties this would be predicted to confer upon tumor cells are sufficient metabolic flexibility to allow for increased TCA cycle-derived anabolic substrate production while avoiding the unsustainable toxicities and TCA cycle-inhibitory properties associated with long-term mitochondrial hyperpolarization and ATP over-production ^21,24^. One way such a state could be achieved would be by activating the mPTP-like function of F_o_ to maintain a normal ΔΨm. Indeed, preliminary studies showed BY1 cells to have higher rates of oxygen consumption relative to ATP production than BY3 cells (Supplementary Figure 6A and B). These results suggest that mitochondria in BY1 cells are less coupled due to enhanced inner membrane leak.

The above dynamic model makes several additional predictions that we tested with appropriate pairwise combinations of WT BY1/BY1-MT-ATP6-c and WT BY3/BY3-TALED cell lines under carefully controlled conditions. Chief among these predictions was that, in its capacity as the mPTP, any increases in free F_o_ would increase the proton leak between the IMS and matrix so as to relieve, limit or prevent mitochondrial hyperpolarization during episodic TCA cycle and ETC hyperfunction ^31,32,75^. The presence of such a transitory mPTP could readily explain the divergence in the rates of ATP synthesis and oxygen consumption between BY1 and BY3 cells (Supplementary Figure 6A and B). Indeed, we found the proton leak of WT BY1 cells to be greater than that of WT BY3 cells. Moreover, this leak was reduced in BY1-Mt-atp6-c cells and increased in BY3-TALED cells (Figure 4A). In agreement with these findings and a second prediction, the ΔΨm of WT BY1 cells was lower than that of BY3 cells (Figure 4B). Neither the partial restoration of Mt-atp6 nor its TALED-mediated mutation significantly altered ΔΨm in either of their respective cell lines. Thus, the ability to simultaneously maintain high levels of TCA cycle function and oxygen consumption is best explained by a proton leak via free F_o_ that, when necessary, allowed partial ΔΨm dissipation without generating excessive ATP via intact CV.

**Figure 4.**
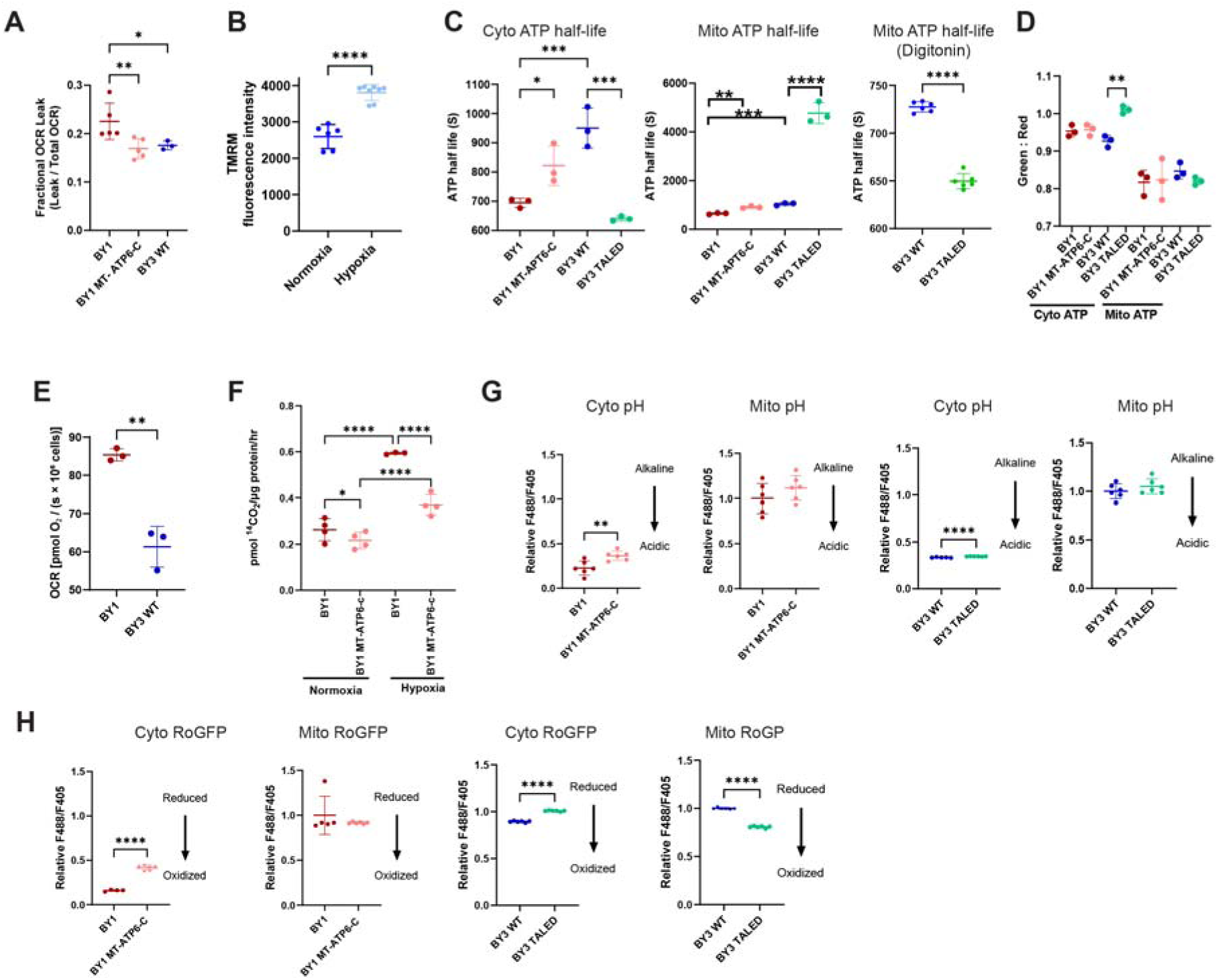
Metabolic properties of BY1 and BY3 cells are altered by F_o_-F_1_ dissociation in ways that support high levels of TCA cycle and ETC function. (A). Free F_o_ increases proton leak. The indicated cell lines, harvested in log-phase growth, were suspended in MiR05 respiration buffer, permeabilized with digitonin and sequentially exposed to cytochrome c, malate, pyruvate, glutamate, succinate, octanoylcarnitine, and glycerophosphate, The OCR at this stage was designated as leak respiration given that no ATP was being produced. ADP was then added to initiate OXPHOS, followed by stepwise titration of carbonyl cyanide chlorophenylhydrazone (m-Cl-CCP) to promiscuously dissipate the proton gradient and achieve maximal uncoupled respiration, thus allowing absolute proton leak to be quantified. (B). ΔΨm is lower in BY1 versus BY3 cells. Cells in log-phase growth were stained with TMRM and average perk fluorescence was assessed by flow cytometry. (C). ATP half-lives are shortened by CV dissociation. The indicated cells, each stably expressing Cyto- or Mito-targeted iATPSnFR2HaloTag ATP sensors, were re-suspended in ice-cold MiR05 buffer lacking glucose (Oroboros, Inc.). After equilibrating to room temperature for 20 min, 2-DG and the CV inhibitor oligomycin were added to a final concentrations of 100 mM and 2.5 μM, respectively and flow cytometric analysis was performed to quantify t_1/2_’s. In other experiments performed in the same manner, BY3 and BY3-TALED cells were permeablized with digitonin and the ANT inhibitor carboxyatractyloside was added to block the export of ATP from the mitochondria and to allow for an assessment of the true intra-mitochondrial ATP t_1/2_. (D). Basal ATP levels are largely maintained in the presence of CV dissociation. The indicated cells, each stably expressing Mito- or Cyto-targeted iATPSnFR2HaloTag ATP, were grown under the indicated conditions, stained with Janelia Fluor JFX650 HaloTag® Ligand and subjected to moving average continuous flow cytometry. (E). OCRs are altered by changes in F_o_-F_1_ association. The indicated digitonin-permeabilized cells were exposed to pyruvate, malate, glutamate, succinate and ADP. OCRs were quantified with an Orboboros respirometer. (F). FAO in WT BY1 cells is suppressed by enforcing MT-ATP6-c expression. FAO was quantified by measuring the release of ^14^CO_2_ from ^14^C-labeled palmitate. (G). Mitochondrial matrix pH of cells with increased free F_o_ is maintained in the normal range. The indicated cell lines, stably expressing Mito- or Cyto-targeted pSypHER GFP were subjected to flow cytometry during log-phase growth ^78^. (H). The mitochondrial matrix of cells with free F_1_ tends to remain relatively oxidized. The indicated cell lines stably expressing Mito- or Cyto-targeted roGFP were subjected to flow cytometry during log-phase growth ^78^.

A third prediction was that free F_1_ in the mitochondrial matrix of BY1 cells would shorten ATP’s half-life (t_1/2_) and prevent its toxic accumulation by virtue of serving as an ATP hydrolase. This would be in agreement with the finding that these cells also synthesize mitochondrial ATP at higher rates and that overall rates of ATP turnover are high (Supplementary Figure 6). Relying upon intact cells that stably expressed Cyto- or Mito-targeted versions of FR2HaloTag ATP sensors ^31,32^, we indeed showed the ATP t_1/2_ in WT BY1 cells to be shorter than that of WT BY3 cells in both cellular compartments and that the t_1/2_‘s were lengthened in BY1-Mt-atp6-c cells (Figure 4C).

The findings with BY3 cells were somewhat more complex. Whereas the cytoplasmic ATP t_1/2_ was indeed shorter in intact BY3-TALED cells, the mitochondrial t_1/2_ was markedly longer. However, it was shorter if the cells were first permeabilized with digitonin, which dissipated ATP in the cytoplasm (Figure 4C). This suggested that cross-talk between the cytoplasm and mitochondria, likely mediated by variations in ATP levels, contributed to the prolongation of mitochondrial ATP half-life. Despite these differences, our findings could be generally reconciled with the fourth prediction, namely that basal levels of ATP would remain normal or even low regardless of the level of free F_1_. Indeed, although some significant differences were noted in cytoplasmic ATP levels, their variation was <10% (Figure 4D).

The fifth prediction was that F_o_-F_1_ dissociation would be associated with more pronounced responses to standard TCA cycle substrates as already shown for tumors (Supplementary Figure 4C) ^41,76^. A corollary to this and thus a sixth prediction was that such cells might also expand their preference for alternative and more energy-rich sources of TCA cycle substrates such as fatty acids ^19^. We therefore utilized Oroboros respirometry to quantify OCR changes in response to pyruvate, malate, glutamate and succinate in each of the 4 cell lines. Indeed, WT BY1 cells demonstrated greater response to these substrates than did WT BY3 cells (Figure 4E). Moreover, the response was reduced in BY1-MT-ATP-c cells and increased in BY3-TALED cells. FAO by WT BY1 cells, as measured by the release of ^14^CO_2_ from ^14^C-labeled palmitate, was also greater relative to BY1-MT-ATP6-c cells and an even more pronounced difference was noted in cells maintained under hypoxic conditions (Figure 4F). Similar experiments performed with WT BY3 and BY3-TALED cells showed FAO rates that were too low to allow for accurate or reproducible comparisons between the groups (not shown).

The seventh prediction was that the more pronounced proton leaks of WT BY1 and BY3-TALED cells (Figure 4A) would acidify their respective mitochondrial matrices. However, because this, like excessive ATP, strongly inhibits the TCA cycle, the protons might be transported to the cytoplasm and/or excreted so as to maintain the alkaline conditions that typify most cancer cells ^22,24,43,77,78^. We thus evaluated mitochondrial matrix and cytoplasmic pH in each of the 4 cell lines using stably-expressed pSypHER reporter vectors ^78^. This showed the mitochondrial matrix pH’s of WT BY1 and BY1-MT-ATP6-c to be indistinguishable whereas the cytoplasmic pH of the former was indeed more acidic (Figure 4G). Similarly, the mitochondrial matrix pH’s of WT BY3 and BY3-TALED cells were identical although the cytoplasmic pH of the latter was more alkaline (Figure 4H).

Oxphos reduces NAD^+^ to NADH, which also strongly inhibits the TCA cycle and ETC ^21,22,24^. Thus, our eighth prediction was that, to avoid this additional source of metabolic suppression, tumor cells with higher levels of F_o_-F_1_ dissociation might actually contain normal or even low mitochondrial NADH:NAD^+^ ratios, with excess electrons being transported to the cytoplasm, likely via a reversal of the malate-aspartate shuttle ^79^. The excess cytoplasmic NADH could then be used to further support the Warburg effect by, for example, serving as a co-factor for the lactate dehydrogenase-mediate conversion of pyruvate to lactate. Thus we utilized mito- and cyto-roGFP to compare the redox states in intact cells ^78^. In keeping with this prediction, the mitochondrial redox states of WT BY1 and BY1-MT-ATP6-c cells were identical whereas the cytoplasm of BY1-MT-ATP6-c cells was somewhat more reduced. Larger and more significant differences were observed between WT BY3 and BY3-TALED cells where, again in keeping with the prediction, the mitochondrial matrix of the latter was more oxidized whereas the cytoplasm was more reduced (Figure 4H). Collectively, these results provide evidence that, in cells with evidence of F_o_-F_1_ dissociation, high levels of TCA cycle function can be sustained by maintaining normal or low intra-mitochondrial concentrations of products that otherwise inhibit the TCA cycle and ETC (Figure 4D,E,F, G and Supplementary Figure 4B and C).

### Inhibiting Mt-atp6 by interfering with mitochondrial DNA integrity or protein synthesis in non-transformed cells promotes CV dissociation

Mt-atp6 and Mt-atp8 are the only mitochondrial-encoded CV subunits that are co-translated and coordinately regulated (Figure 1D and Supplementary Figure 3) ^80^. We thus reasoned that interfering with their synthesis in non-transformed cells would impact F_o_-F_1_ dissociation in ways that at least partially mimic what has been observed in primary tumors, their derivative cell lines and in Leigh syndrome (Figures 1A and B, 2A and B and Supplementary Figure 1A-C) ^36,37,39^. This would provide independent evidence that maintaining proper stoichiometric balances among these proteins were essential to CV’s structural integrity, particularly given that interfering with mitochondrial-specific gene expression can impact nuclear-encoded mitochondrial-specific genes (Figures 1A,B,G and H; 2A and B and Supplementary Figure 5C and D) ^81^. To test this, we inhibited Mt-atp6 in NIH3T3 murine fibroblasts in 3 different ways. First, we generated “Rho0” cells in which mtDNA is gradually depleted with low concentrations of ethidium bromide (EtBr) ^82,83^. Second, we blocked mitochondrial protein synthesis with doxycycline, which binds to and inhibits the mitochondrial 30S ribosomal subunit ^84^. Finally, we again blocked mitochondrial protein synthesis but used chloramphenicol, which binds the 23S rRNA component of the 50S ribosomal subunit and inhibits its peptidyl transferase activity ^85^. In the first case, EtBr exposure caused a dose-dependent and ultimately near-total loss of intact CV and a marked increase in free F_1_ (Figure 5A). Following a 2 wk recovery period in EtBr-free medium, free F_1_ was no longer detected and intact CV was restored. Both findings paralleled changes in mtDNA and Mt-atp6 protein content although only partial restoration of the former was needed to achieve full recovery of the latter (Figure 5B and C). Moreover, changes in the relative amounts of Mt-atp6 and nuclear-encoded Atp5f1a over the course of the study spoke to the previously mentioned stoichiometric imbalance and complex regulatory relationships that exist between mitochondrial- and cytoplasmically-encoded components of the respiratory chain (Figure 5C, Supplementary Figure 2) ^86^. Doxycycline and chloramphenicol also resulted in time-dependent declines in Mt-atp6 levels, albeit with somewhat different kinetics and degrees of depletion (Figure 5D). *In situ* ATPase assays performed 48 hr after exposure to the antibiotics showed increases in free F_1_ in both cases, although these were not as pronounced as they were with EtBr (Figure 5E). Cells expressing the Mt-atp6-c fusion protein were less susceptible to free F_1_ generation in response to EtBr and doxycycline (Figure 5F-H). Finally, and in keeping with previous findings that free F_1_ accelerates ATP hydrolysis (Figure 4C), we noted significant reductions in ATP half-lives in BY3 cells following exposure to hypoxia or chloramphenicol (Figure 5H). Collectively, these and our foregoing studies established that the depletion of Mt-atp6 via processes involving transformation, *Mt-atp6* gene mutagenesis or mitochondrial DNA depletion are all associated with the loss of CV integrity and reversible F_o_-F_1_ dissociation.

**Figure 5.**
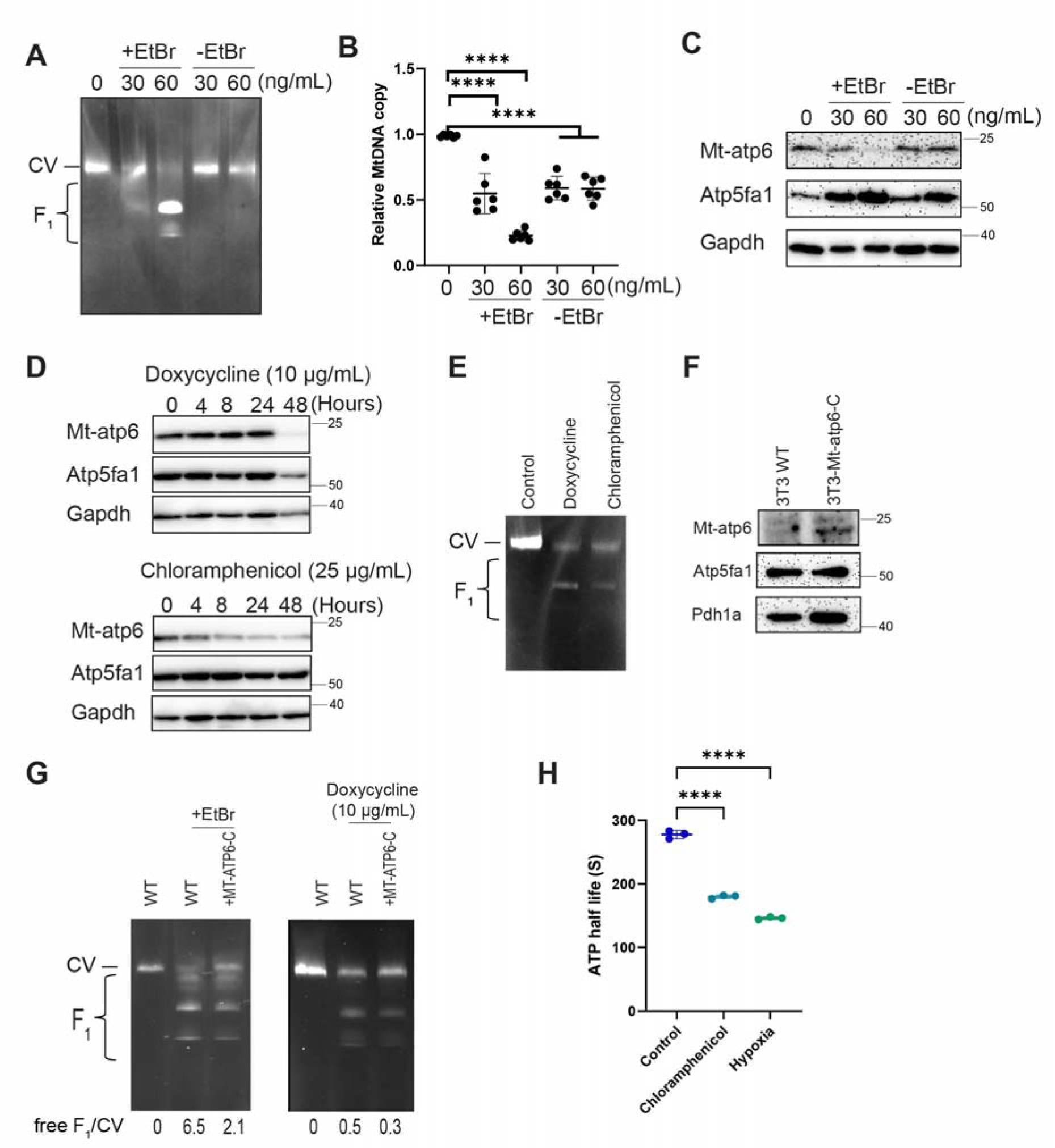
Depletion of mtDNA or inhibition of mitochondrial protein synthesis promotes F_o_-F_1_ dissociation in ways that can be altered by genetically manipulating Mt-Atp6 levels. (A). Appearance of free F_1_ in NIH3T3 fibroblasts during Rho0 cell generation. Cells were maintained in 30 or 60 ng/ml of EtBr for 2 days (lanes 2 & 3) and then switched to EtBr-free medium for 2 wks to allow for recovery (lanes 4&5). At each point, *in situ* ATPase assays were performed. (B). mtDNA content of EtBr-treated cells described in (A) and normalized to that of the nuclear-encoded *Mlx* gene ^43^. (C). Immuno-blot showing that mitochondrial-encoded Mt-atp6 but not nuclear-encoded Atp5f1a α subunit is depleted in a dose-dependent manner by EtBr (+EtBr) and then restored following subsequent recovery in EtBr-free medium (-EtBr). (D). Immunoblot showing time-dependent Mt-atp6 depletion in NIH3T3 cells in response to doxycycline or chloramphenicol treatment. (E). *In situ* ATPase assays performed on NIH3T3 cells exposed to doxycycline (10 μg/ml) or chloramphenicol (25 μg/ml) for 48 hr. (F). Stable expression and mitochondrial localization of Mt-atp6-c in NIH3T3 cells. Mitochondria from control (WT) cells or those stably expressing the Mt-atp6-c fusion protein were purified as described in Figure 2H. Loading control immuno-blots were performed for the mitochondrial-localized proteins Pdha1a and ATP5fa1. (G). Identification of TALED mutations in NIH3T3 cells using the T7 endonuclease assay described in Figure 2I. (H). Enforcing Mt-atp6-c or mutating endogenous Mt-atp6 with TALEDs, respectively, alters NIH3T3 cell tsusceptibility to EtBr-mediated F_o_-F_1_ dissociation. *In situ* ATPase assays were performed on control, untreated NIH3T3 cells or on cells cultured in EtBr (30 ng/ml) as described in (A). (I) Hypoxia and chloramphenicol reduce ATP t_1/2_ in BY3 cells. Cells stably expression Mito-targeted iATPSnFR2HaloTag were exposed to 1% oxygen or 25 mg/ml chloramphenicol for 48 hr. They were then stained with Janelia Fluor JFX650 HaloTag® Ligand and subjected to moving average continuous flow cytometry as described in Figure 4C.

## Discussion

Previous studies have suggested that Complex V’s dissociation into its component F_o_ (mPTP) and F_1_ (ATPase) functional domains is not necessarily detrimental and may even be advantageous, particularly if it is reversible ^31,32^. When it is not, due to the *MT-ATP6* mutations associated with Leigh syndrome, affected individuals typically display varying degrees of cardiomyopathy, neuropathy, diabetes mellitus and lactic acidosis ^26,33,35,53,87^ ^26,35,36,54,88^. Their mitochondria can show high levels of free F_1_ ATPase, decreased rates of ATP synthesis, increased proton leak and ΔΨs that are either higher or lower than normal ^26,33,35,53,87^. The variability of these clinical and metabolic findings may reflect the severity of different mutations, the degree of heteroplasmy, the identities of the tissues being studied and/or different metabolic states of the cell at the time of examination. It has been demonstrated that restoring CV structural integrity via the delivery of mitochondrial-targeted wild-type MT-ATP6 can variably rescue the metabolic dysfunction ^33,51,52,66^. Collectively, these findings indicate that MT-ATP6’s regulation of proton flow through F_o_ is directly related to its ability to contribute to mPTP function and to stabilize the F_o_-F_1_ interaction by securing the peripheral stalk subunit OSCP and other accessory proteins to F_o_ and the IMM (Figures 2C and& F) ^27,29^.

Low levels of otherwise wild-type MT-ATP6 (and co-translated MT-ATP8) are common features of several murine and human tumor types, including HBs, HCCs and renal cancer where free F_1_ ATPase activity has also been noted (Figure 1C and D and Supplementary Figures 2 and 3) ^74^. Moreover, and specifically in the case of genetically-defined HBs and HCCs, unequal declines in transcripts encoding the CV’s subunits may contribute to stoichiometric imbalances that could underlie the differential prominence of F_o_-F_1_ dissociation (Figures 1C-G and 2B and Supplementary Figures 3D and E and 4). Functionally, these quantitative differences would behave similarly to the qualitative one associated with Leigh syndrome and other CV genetic disorders but offer the advantage of being reversible and potentially susceptible to both qualitative and quantitative forms of regulation ^89^. Thus, our work leaves open the possibility that other CV subunits may be involved in this process, either individually or cooperatively. For example, of the 22 human tumor types for which sufficient numbers of adjacent normal tissues could be evaluated, the most frequently down-regulated gene after *MT-ATP6* and/or *MT-ATP8* was *ATP5PF* that was seen in 8 tumor types (Figure 1D)*. ATP5PF*, which encodes the peripheral stalk protein F6 that also maintains the F_o_-F_1_ association via OSCP, has been reported to be down-regulated in individuals with Alzheimer’s disease who display increases in free F_1_ ^29,49^. Elsewhere, *ATP50,* which encodes OSCP itself, is sometimes down-regulated in several other tumor types (Figure 1D). Collectively, these findings indicate that Complex V dissociation may occur under a variety of circumstances and disease states and be mediated by stoichiometric imbalances among proteins other than MT-ATP6 and MT-ATP8, all of which are nuclear-encoded. This would potentially allow CV integrity to be subject to non-mutually exclusive forms of both mitochondrial and extramitochondrial control in response to changes in various environmental influences that would afford high levels of flexibility ^26,29,49,86,90^.

Several features distinguish the cancer-related CV defects described here from those of Leigh syndrome. In the latter case, missense or truncation mutations in MT-ATP6, and less commonly MT-ATP8, irreversibly inactivate the respective protein, in turn restricting the degree to which CV dissociation can be reversed or otherwise regulated. The ensuing abnormalities in Oxphos plus inadequate compensation via the Warburg effect, likely account for the failure to coordinate the production and maintenance of normal levels of ATP ^36,37,39,53^. *MT-ATP6* mutations are not only rare in human cancer but seldom involve the amino acids associated with Leigh syndrome ^91^. Quantitative changes of the type that we have described are not only more likely to cause stochastic imbalances among CV subunits but are more amenable to correction in response to altered environmental or proliferative demands (Figures 1B-D, 2B, 3B and Supplementary Figure 3). The fact that not all HBs or cell lines derived from them express free F_1_, despite having otherwise identical genetic backgrounds, may reflect subtle differences in this CV subunit imbalance and/or total mitochondrial mass (Figure 1C and D, Supplementary Figures 1C and 4A,D and E). The metabolic flexibility of cancer cells that allows them to engage in compensatory Warburg-type respiration ^2,13,15^ also likely provides further protection against any impairments in mitochondrial ATP supplies arising as a result of its reduced synthesis or increased F_1_-mediated hydrolysis. The Warburg effect, coupled with fundamental differences in how F_o-_F_1_ dissociation is achieved and controlled, may thus turn Leigh syndrome-like liabilities into assets in cancer.

Previously reported difficulties in reconstituting detectable levels of human MT-ATP6 are likely due to the protein’s high hydrophobicity, its short half-life and its inefficient mitochondrial targeting when ectopically expressed in the nucleus ^33,51,52^. We encountered similar problems in stably expressing murine Mt-atp6 despite employing several combinations of vectors, mitochondrial targeting signals and epitope tags (not shown). Ultimately, we achieved consistent and stable expression in both transformed and non-transformed cells as well as proper mitochondrial localization when Mt-atp6 was expressed as a fusion protein in association with the c subunit (Figure 2F-H). This allowed for F_o_-F_1_ re-association and corrected many of the abnormalities associated with free F_o_ in BY1 cells (Supplementary Figure 5). It seems improbable that the c fusion addition itself performed any function other than to stabilize Mt-atp6 and guide its incorporation into CV given that doing so would likely impede c ring rotation and exacerbate rather than correct the proton leak (Figure 2F) ^26,27,29^.

While TALED-mediated dissociation of CV into its component F_o_ and F_1_ domains in BY3 cells provided results that complimented those generated in BY1-MT-ATP6-c cells, they too were nevertheless imperfect. This was likely due to the mutations introduced into the *Mt-atp6* gene being identical to those from individuals with Leigh syndrome ^36,37,39,53^. As a result, while further increases in free F_1_ could be achieved by factors such as hypoxia, pre-existing CV defects of BY3-TALED cells were not reversible and were thus less responsive to certain environmental cues (Figure 3A). This limited metabolic flexibility may thus explain why Leigh syndrome-like MT-ATP6 mutations are rare in cancer cells, which instead prefer the greater adaptability afforded by quantitative changes. Despite the less than ideal means of restoring and inhibiting *Mt-atp6* expression, our finding were nevertheless in surprising agreement with our proposed model of anabolic substrate and ATP balancing in cancer (Figure 6).

**Figure 6.**
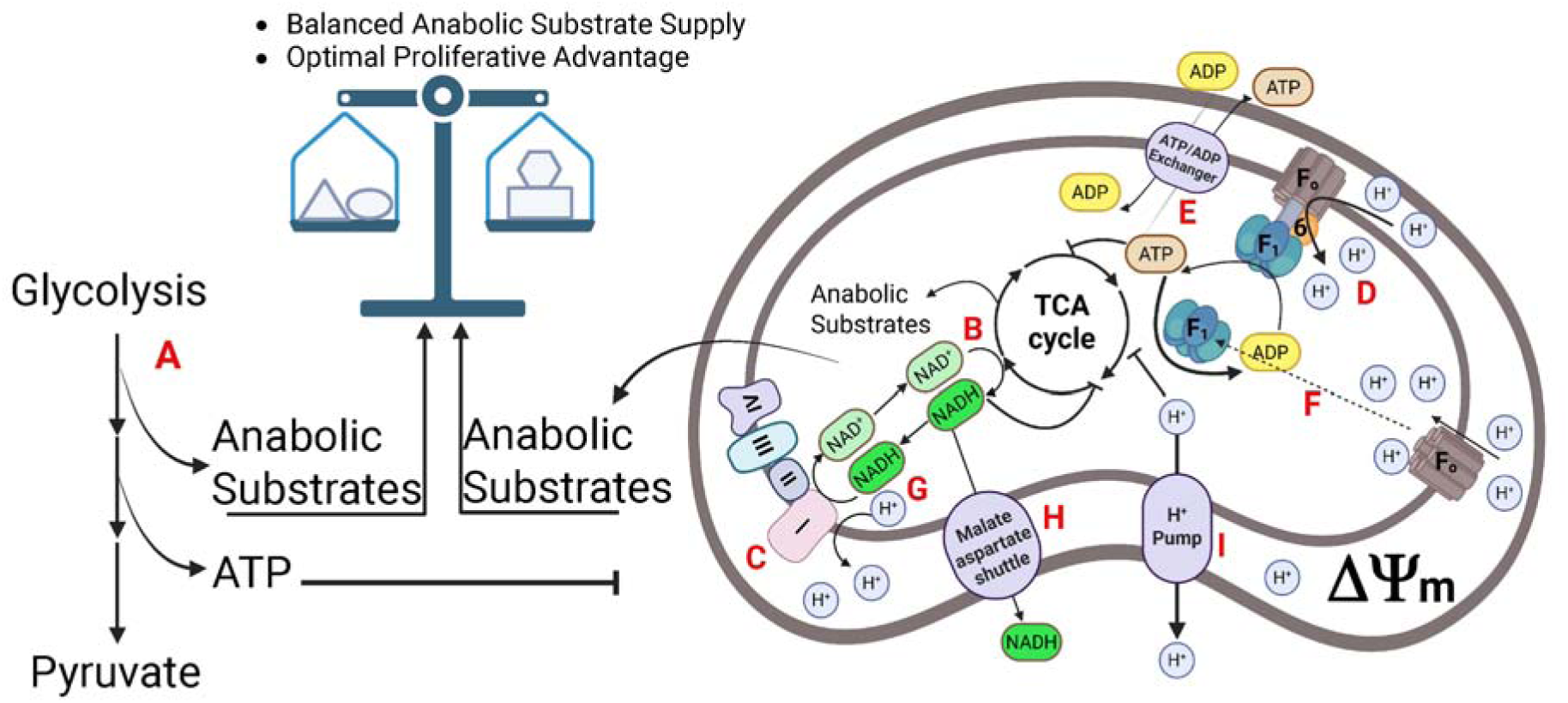
A model that balances the production of glycolysis- and TCA cycle-derived anabolic substrates. (**A**). Increased glycolysis during Warburg effect-type respiration increases the supply of glucose-derived ATP and anabolic substrates. The former will tend to suppress mitochondrial ATP production to avoid the toxicity associated with ATP excess whereas the latter will result in anabolic substrate imbalance. To achieve optimal tumor growth rates, the supplies of anabolic substrates and ATP must be balanced with those derived from TCA cycle precursors and Oxphos. (**B**) The TCA cycle donates electrons to the ETC using NADH as the door (dark green) and NAD^+^ as the acceptor (light green). (**C**). Energy generated by the ETC pumps protons into the IMS and establishes the ΔΨm. (D) Controlled proton return to the matrix via CV’s F_o_ domain provides the electromotive force to generate ATP via F_1_. MT-ATP6 contributes to the formation of the proton channel (Figure 2F). (E) ATP is transported to the cytoplasm via the ATP/ADP carrier, thereby reducing its inhibitory effect on the TCA cycle and ETC. (F) In MT-ATP6’s absence, some CV dissociates into F_o_ and F_1_. Free F_o_, now serving as the mPTP, allows proton return to the matrix without generating ATP while free F_1_ acts as an ATPase in a manner that is independent of F_o_. This represents the key processes by which physiologic ΔΨm and ATP levels are maintained and regulated in the face of high TCA cycle function. (G) Excess NADH, also a potent inhibitor of the TCA cycle and the ETC, is exchanged for NAD^+^ via reversal of the malate-aspartate shuttle thereby maintaining low concentrations (H). (**I)** Excess matrix protons, generated by CV and free F_o_, and also inhibitory of the TCA cycle and ETC, are transferred to the cytoplasm.

Despite the fact that TALED-generated murine *Mt-atp6* mutations did not serve as ideal substitutes for mimicking the F_o_-F_1_ dissociation that occurs in cancer, they nevertheless underwent considerable *in vivo* and *in vitro* selection following their initial generation (Figure 3E). This serves as direct evidence that certain Leigh syndrome-associated mutations or mutational combinations are both positively and negatively selected in ways that ultimately contribute to the target cell’s proliferative advantage. This almost certainly reflects the fact that different *Mt-atp6* mutant proteins retain variable amounts of residual function and thus may differentially contribute to the retention of CV’s stability in ways that favor or disfavor its reversible dissociation in a transformed environment. The identities of these mutations and their relative levels of expression may also reflect the degree to which individual tumors are capable of modulating the Warburg effect, which can vary considerably among individual tumors, even when generated by identical oncogenic drivers ^43,92^.

Based on immuno-blotting, the actual amount of F_o_-F_1_ dissociation in tumor cells appears to be considerably greater than that suggested by *in situ* ATPase assays (Figures 1A and B, 3A and Supplementary Figures 1 and 5). This could reflect several non-mutually exclusive factors, including differences in the intrinsic enzymatic activities of various F1 states, a reduced ability of the α subunit to enhance the intrinsic ATPase activity of the β subunit due to subtle conformational changes, the lack or alteration of the γ subunit that stimulates the intrinsic ATPase activity of the core α_3_β_3_ core F_1_ ATPase and inherent limitations of the *in situ* ATPase assay itself (Figure 2C) ^26,93,94^. In non-transformed NIH3T3 cells, the fact that virtually all intact CV-associated ATPase activity is lost following EtBr treatment (Figure 5B) indicates that even near-total F_o_-F_1_ dissociation is well-tolerated and reversible, at least over the short term. The lesser amounts of reversible dissociation and reassociation seen in cancers may be sufficient to permit recovery from transient episodes of mitochondrial membrane hyper- or hypo-polarization, prevent unsustainable swings in energy production and avert what would otherwise be substantial growth impairments and/or toxicities while fulfilling its role in regulating the supply of TCA cycle-derived anabolic substrates. Such levels also appear to be sufficiently low so as to avoid impairing overall mitochondrial integrity. The above-proposed regulatory role further suggests that free F_o_ and F_1_ are likely distributed in small amounts among all mitochondria rather than being concentrated among a selective few where they would likely prove more deleterious than beneficial.

Exposure of non-transformed cells to EtBr, doxycycline and chloramphenicol led to F_o_-F_1_ dissociation via different mechanisms that ultimately converged on *Mt-atp6* in ways that are consistent with our observations in primary tumors and their derivative cell lines (Figure 5A-E). The consequences of EtBr, which ultimately generates cells lacking mitochondria and mtDNA, are more akin to what is observed in transformed cells where numerous mitochondrial genes encoded by both the mtDNA and nuclear genomes are deregulated and create stochastic imbalances among CV subunits and F_o_-F_1_ instability (Supplementary Figures 2 and 4D-G) ^82,83^. Doxycycline and chloramphenicol differ from EtBr in that they more specifically target the expression of mitochondrial-encoded proteins, only 2 of which (Mt-atp6 and Mt-atp8) are subunits of CV ^84,85^.

The shorter ATP t_1/2_ in the mitochondrial matrix of cells with free F_1_ likely reflects, at least in part, the CV-independent intrinsic ATPase activity of this domain (Figure 4C). Since this activity is not subject to the F_o_-mediated control associated with intact CV, it may well be regulated by other factors that prevent the promiscuous depletion of ATP, while maintaining concentrations that are sufficient to support anabolic needs without inhibiting TCA cycle function. Among the most obvious ways this control could be exerted would be at the level of α and β subunit interaction, which greatly enhances the intrinsic ATPase activity of the latter subunit as mentioned above ^94^. Interactions between the β subunit and Atp5if1/If1, a potent inhibitor of the former’s intrinsic ATPase activity, as well as various metabolites generated during the course of high-level TCA cycle activity, represent additional means of exerting such regulation that are deserving of further investigation ^63,93^. Importantly in cells with free F_1_, the shorter t_1/2_ of ATP (Figure 4C) may reflect not only its rate of hydrolysis but also its transport from the mitochondria to the cytoplasm, enzymatic contributions from non-F_1_ ATP hydrolases such as the V- and P-type ATPases that are often elevated in cancers and differences in growth ^95,96^. Regardless of the basis for the shortened t_1/2_, the ultimate effect should be one that contributes to TCA cycle activity and the provision of its unique anabolic substrates.

Our model resolves some of the seemingly paradoxical aspects of cancer cell metabolic re-programming (Figure 6) ^2,13,15,17^. First, it establishes that the loss of mitochondrial mass observed in most tumors can be offset by a small but more efficient subpopulation of mitochondria (Supplementary Figure 4A) ^14,95,97^. Energy that would normally be expended in supporting a large mitochondrial mass can instead be directed toward growth and proliferation. In the remaining more active mitochondria, maintaining normal ΔΨm and ATP levels as a result of the partial uncoupling of the TCA cycle and ETC from ATP synthesis by CV allows the supply of anabolic substrates to be increased as necessary to match those supplied by the high rates of Warburg-type glycolysis. These mitochondria are also able to select from a wider range of fuel sources and to become more reliant on anaplerosis to maintain supplies of TCA cycle intermediates, thus explaining other metabolic changes often associated with cancers (Figure 4F) ^98–100^. The rapid reversibility of CV dissociation provides the necessary metabolic flexibility to allow a return to normal or even lower rates of TCA cycle function during times of reduced proliferative activity, reduced anabolic substrate demand or nutrient deprivation.

The findings presented here raise several questions that will be important to address in future work. These include whether factors other than quantitative alterations of CV subunits impact Mt-atp6’s stabilization of the Fo-F_1_ association. The most obvious candidates to effect such changes would be molecules whose concentrations within the mitochondrial matrix reflect the activity of the TCA cycle and/or the organelle’s energy status. These could include TCA cycle substrates, ADP, ATP, NAD^+^ and NADH. Their interaction with Mt-atp6 might be expected to involve hydrophilic regions of the protein that are exposed to the matrix rather than the hydrophobic and IMM-embedded regions within which Leigh syndrome mutations reside ^35–37,39,50,68^. It is also possible that the effects on Mt-atp6 are indirect and occur as a result of interactions with other CV subunits such as Mt-atp8, the c subunit or OSCP ^28,29^. Another important question concerns the means by which known inhibitors of the TCA cycle and ETC such as protons, ATP and NADH are maintained at low levels within the matrix when free F_1_ is present despite high rates of TCA cycle function. Although we have not explored in detail how these low levels are achieved or maintained, our findings suggest that they are likely to be important for regulating not only the extent of CV dissociation but the degree to which its component F_o_ and F_1_ domains maintain maximal function once this occurs.

## Materials and Methods

### Animals and tumor generation

All animal studies were approved by the University of Pittsburgh Institutional Animal Care and Use Committee (IACUC). Hepatoblastomas (HBs) were generated as previously described in 4-6 wk old FVB/N mice ^41–44^. Two Sleeping Beauty (SB) vectors were used for this and were delivered via hydrodynamic tail vein injection (HDTVI). The first vector encoded a human HB-derived mutant form of β-catenin bearing a 90 amino acid in-frame deletion (Δ90, hereafter “B”) and the second encoded yes-associated protein (YAP) bearing an oncogenic missense mutation (YAP^S127A^, hereafter “Y”) ^42^. Both B and Y constitutively localize to nuclei and generate “BY” tumors with near 100% efficiency when co-expressed but no tumors when expressed individually ^42–44^. These plasmids, along with an additional non-SB vector encoding a SB transpose were purified using Qiagen Maxi Plasmid Isolation Kits (Qiagen, Inc. Germantown, MD). 10 μg of each SB plasmid and 2 μg of the SB transpose-encoding vector were delivered by HDTVI over ∼3-9 sec in a total volume of 2 ml of in 0.9% saline solution. For some studies, SB vectors encoding other patient-derived β-catenin mutations were used to generate HBs using the same approach ^43^.

FVB/N-Tg(tetO-MYC)36aBop/J and LAP-tTA mice (B6.Cg-Tg[Cebpb-tTA]5Bjd/J) were obtained from Jackson Laboratories (Bar Harbor, ME) and crossed as previously described, with the offspring hereafter being referred to hereafter as Tet-*MYC* mice ^45^ Animals were provided standard mouse chow *ad libitum* as well as drinking water that contained doxycycline (100 μg/ml). HCCs were induced by removing the doxycycline from the water. Control livers were obtained from mice that had been maintained on doxycycline (d0) or from which doxycycline had been discontinued for 3 days or 7 days but in which tumors had yet to develop (d3 and d7, respectively) ^45^. ∼30 days after discontinuing doxycycline, large HCCs were removed as were additional tumors 3 days or 7 days after reinstating doxycycline and inducing regression (R3 and R7). Recurrent tumors (RT) were induced in mice in which initial tumors had been allowed to completely regress for 2-3 months.

### Generation of Mt-atp6 and Mt-atp8 expression vectors

To enable ectopic expression of mitochondrial genes in mammalian cells, codon-optimized constructs for human *MT-ATP6* and murine *Mt-atp8* were generated (Gen Script Biotech, Inc. Piscataway, NJ). Codon-optimized *MT-ATP6*, fused at its 5′ end to the mitochondrial targeting signal (MTS) of human *ATP5MC1*, was PCR-amplified from the pCMV6-G1-ATP6-Myc-FLAG plasmid (Addgene #86851). Codon-optimized *Mt-atp8* was synthesized *de novo*. Full sequences are provided in Supplementary File 1. To enhance mitochondrial import and facilitate detection, the mature coding region of murine *Atp5mc1* (encoding the ATP synthase c subunit) was PCR-amplified from reverse-transcribed total mouse liver RNA. For *Mt-atp6*, a fusion construct was generated with the *ATP5MC1* MTS at the N-terminus, followed by *Mt-atp6*, and the mature *Atp5mc1* cDNA fused in-frame with a C-terminal V5 epitope tag. For *Mt-atp8*, the coding sequence was also fused to the *ATP5MC1* MTS at the 5′ end and a 3×FLAG epitope at the 3′ end. All constructs were cloned into the pSBbi-RP Sleeping Beauty (SB) transposon-based expression vector (Addgene #60513), which includes a dTomato reporter and puromycin resistance cassette. Constructs were fully sequenced (Genewiz/Azenta Life Sciences, Burlington, MA) to confirm sequence accuracy and proper orientation. For stable transfections, 2 μg of each SB vector plus 0.2 μg of a non-SB vector encoding SB transposase ^41,43,44^ were mixed and introduced into BY1 or BY3 cells that had been seeded the previous day in 6 well plates so as to be at 30-50% confluence at the time of transfection. Transfections were performed with Lipofectamine-2000 according to the instructions provided by the vendor (Thermo Fisher, Pittsburgh, PA).

### Generation and characterization of TALED-mediated mutations in *Mt-atp6*

The plasmids TALED_Left-mouse-ATP6-139N and TALED_Right-mouse-ATP6-1397C-AD(V28) (hereafter “TALED vectors”) ^67^ were obtained from Addgene, Inc. (Watertown, MA). They were transiently co-transfected into BY3 or NIH3T3 murine fibroblasts (obtained from the ATCC, Manassas, VA) at a 10:1 ratio with the Cyto- and Mito-targeted roGFP or pSypHER plasmids described above followed by FACS purification and G418-selection of the GFP+ population 2 days later. To evaluate the efficiency of TALED-mediated mutagenesis, a 504 bp fragment of murine mitochondrial DNA (mtDNA) spanning the *Mt-atp6/8* gene was amplified with Platinum SuperFi DNA polymerase (Thermo Fisher) from 10 ng of total cellular DNA in a 50 μL PCR reaction using the following primers ^67^: FWD: 5’-CACATACATTTACACCTACTACCC-3’ and REV: 5’- GTTAGAAGGAGGGCTGAAAAGG-3’. 35 cycles of amplification were performed under the following conditions: denaturation: 5 sec x 98C, annealing: 10 sec x 55 C, 20 sec x 72 C followed by a final extension at 72 C x 2 min. To establish the extent of heteroplasmy, the frequency of induced mutations was initially estimated using a T7 endonuclease assay according to directions of the vendor (New England Biolabs, Inc. Ipswich, MA) (Wang). In BY3 cells, individual mutations were then quantified by subjecting the same PCR fragments to next generation high-throughput DNA sequencing (Genewiz).

### Generation of HB cell lines

A detailed description of immortalized HB cell line generation derived from BY tumors is provided elsewhere ^61,101^. Briefly, HBs were induced as described above except that the inocula contained 2 additional non-SB pDG458 Crispr/Cas9 vectors (Addgene, Inc. Watertown, MA), each of which encoded 2 gRNAs directed against exon 2 of the murine *Cdkn2a* locus, which encodes both the p16^INK4a^ and p19^arf^ tumor suppressors in overlapping reading frames. The 2 gRNA sequences for pDG458-Vector 1+2 were 5’-CGGTGCAGATTCGAACTGCGAGG-3’ and 5’-GTCGTGCACCGGGCGGGAGAAGG-3’. gRNA sequences for pDG458-Vector 3+4 were 5’-CTTGGGCCAAGTCGAGCGGCAGG-3’ and 5’-TGCGATATTTGCGTTCCGC TGGG-3’. Multiple ∼1 mm tumor fragments were prepared from several grams of freshly isolated primary tumors tumor, washed in PBS and digested in 0.1% trypsin (Sigma-Aldrich, Inc. St Louis, MO) for 30 min at 37C. The digested tissues were then vigorously vortexed and pipetted several times and incubated in standard tissue culture plates with D-MEM containing 10% fetal bovine serum (FBS), 0.2 mM L-glutamine, 100 units/ml of penicillin G and 100 μg/ml streptomycin that was changed every 3-4 days. Over the next several wks, cells originating from the tumor fragments attached to the surface of the plates and eventually formed dense colonies comprised of proliferating cells atop a monolayer of quiescent cells. These were removed by trypsinization, pooled and expanded for 3-4 additional passages before being stored in liquid nitrogen. Successful inactivation of the *Cdkn2a* locus was confirmed by PCR and high-throughput sequencing and by demonstrating the absence of WT p19^arf^ and p16^INK4a^ proteins as described ^61,101^. For the current study, we relied upon 2 of the original 8 cell lines that were derived: BY1 cells, which expressed free F_1_ under normal conditions of growth and BY3 cells, which did not.

### Subcutaneous (subq) tumor growth studies

For some studies, FVB/N mice were used to compare the *in vivo* growth rates of BY1 and BY3 cell HB cell lines. For these studies, 10^6^ cells of the indicate cell line were injected subq into the flanks of the animals. Tumor dimensions were calculated with calipers on a regular schedule and volumes were determined using the formula V = (l x h x w) x (π/6).

In other studies, nu/nu mice were obtained from Jackson Labs and were used to determine whether WT BY1 cells displayed an *in vivo* growth advantage relative to BY1-Mt-atp6-c cells. Each cell line was stably transfected with the cyto-roGFP expression vector described elsewhere in this section. These cells were then combined with an equal number of untagged cells of the other line (10^6^ each) and subq tumors were generated in 4-6 wk old mice. The 2 groups thus comprised WT BY1 + BY1-M-ATP6^cyto-roGFP^ and WT BY1^cto-roGFP^ + BY1-Mtatp6-c populations that were allowed to compete during the 5-6 wk course of tumor propagation ^61,101^. DNAs from tumors and the input cell populations were then purified and used for TaqMan reactions that quantified the relative amounts of GFP and human β-catenin (derived from B) in a manner that allowed only tumor cell populations to be evaluated. The primers and TaqMan probes used were as follows: GFP-Forward: 5’-AGTGCTTCAGCCGCTACC -3’; GFP-Reverse: GAAGATGGTGCGCTCCTG; GFP TaqMan probe: 5 ′ /56-FAM/TTCAAGTCC/ZEN/GCCATGCCCGAA/3IABkFQ/-3 ′; β-catenin-Forward: 5′-CAGAGTGCTGAAGGTGCTATC-3′; β-catenin-Reverse(region of the SB vector just downstream of the β-catenin insertion site): 5′-GATCTGTCAGGTGAAGTCCTAAAG-3′; β-catenin TaqMan probe: 5′-6-FAM/CCGGCTATTGTAGAAGCTGGTGGAA/3′ Iowa Black® FQ/-3′. Each PCR reaction, which was performed in triplicate, contained 100 ng of input DNA, employed Taq DNA polymerase (NEB, Cat# M0273) and was subjected to 40 rounds of amplification using the following conditions: 95 °C for 10 s; 40 cycles at 95 °C for 15 s, and 60 °C for 1 min. All assays were performed on a CFX96 Touch^TM^ real-time PCR detection system (Bio-Rad). Standard curves from cell populations containing known amounts of each of each cell line were used to ascertain the amounts of each population present in the tumors obtained at the termination of the study.

### Measurements of cytoplasmic and mitochondrial redox state and pH

Plasmids encoding cytoplasmic (“Cyto”) and mitochondrial (“Mito”) targeted roGFP and pSypHer^78^ were introduced into BY1 and BY3 cell lines using SuperFect 2000. For roGFP, GFP-positive cells were sorted by flow cytometry due to the lack of a selection marker. The pSypHer gene, PCR-amplified from the original vector ^78^ using SfiI-adapted primers for both Cyto and Mito versions, was recloned into the pSBbi-Neo vector at SfiI sites and stably selected with 500 μg/ml G418 (Geneticin, Thermo Fisher). Cells were pooled for all subsequent studies. Redox states were measured on cells in early-mid log-phase growth 2 days after re-plating. For both sets of cell lines, flow cytometry was used to measure either the redox state or pH of individual cellular compartments by determining the ratio of 530 nm emission signal intensities upon excitation at 488 nm and 405 nm.^78^.

### Determination of mitochondrial mass, glucose uptake and ΔΨm

Mitochondrial mass was determined by staining cells *in vitro* with nonyl acridine orange (NAO) or Mitotracker Green as previously described ^78^. To assess mitochondrial mass in primary HBs and livers, we relied on a TaqMan-based assay to quantify mitochondrial DNA (mtDNA) as described previously ^43,78^. Briefly, 2 sets of PCR primers were utilized for the assay, with the first designed to amplify a 101 bp region of the mitochondrial cytochrome c oxidase gene and the second designed to amplify a 90 bp region from the D-loop region. PCR reactions were normalized to the TaqMan signal obtained from the amplification of the control apolipoprotein B nuclear-encoded gene. All reactions were performed on a CFX96 Touch^TM^ Real-Time PCR Detection System (Bio-Rad, Inc., Hercules, CA) (95C x 10 sec., followed by 40 cycles at 95C x 15 sec. and 60C x 60 sec.).

Glucose uptake was measured by staining live cells with the fluorescent glucose analog 2-(7-Nitro-2,1,3-benzoxadiazol-4-yl)-D-glucosamine (2-NBDG) and ΔΨm was determined by staining with tetramethylrhodamine methyl ester (TMRM) (both from Thermo Fisher, Inc.). Staining procedures followed the instructions provided by the vendors and results were evaluated by flow cytometry as previously described ^47^.

### Determination of cytoplasmic and mitochondrial ATP content and ATP half-lives

To determine basal ATP levels, cells were stably transfected with either cytosolic or mitochondria-targeted iATPSnFR2-HaloTag sensors (Addgene). These constructs were cloned from pAAV.CAG.(mito). iATPSnFR2.A95A.A119L.HaloTag (Addgene plasmid #209722) using SfiI site-adapted primers, either with or without a 3×COX8 mitochondrial targeting signal. The resulting PCR products were directly cloned into the pSBbi-Neo vector at SfiI sites (Addgene plasmid #60525). ^102^. Cells in log-phase growth were harvested by trypsinization and resuspended in complete DMEM medium. Janelia Fluor® JFX650 HaloTag® Ligand (Promega, Inc., Madison, WI) was then added to the suspension at a final concentration of 200 nM, followed by incubation at 37°C for 30 minutes to allow for HaloTag labeling. After incubation, the staining medium was removed, cells were washed and resuspended in fresh complete growth medium. Cells were then analyzed on a BD Fortessa flow cytometer, with the following settings: JFX650: Excitation 627–640 nm, Emission 670 ± 10 nm; ATPsensor Green: Detected using standard FITC/GFP or relevant green fluorescence channel (excitation/emission around 488/530±30 nm).

To measure mitochondrial ATP half-life, 2–3 x 10^6^ cells stably expressing the above-described Mito-targeted iATPSnFR2HaloTag ATP sensor were harvested by trypsinization and resuspended in 1 mL of Mir05 respiration buffer that contained no glucose (Oroboros, Inc.). Cells were then permeabilized by adding digitonin to a final concentration of 20 µg/mL and incubated for 10 minutes at room temperature. TCA cycle substrates were sequentially added to the cell suspension in the following order and at the indicated final concentrations: ADP (2.5 mM), malate (2 mM) and pyruvate (5 mM). This was followed by the addition of the ANT inhibitor Carboxyatractyloside (5 μM) (Sigma-Aldrich) to block the export of ATP. Finally, oligomycin (20 nM) (Sigma-Aldrich) was added to inhibit the *de novo* synthesis of ATP by CV. Cells were then immediately subjected to continuous flow cytometric analysis at a rate of >400 cells/sec to monitor mitochondrial ATP levels over time.

### Quantification of oxygen consumption rates (OCRs), proton leak and rates of ATP synthesis

OCRs were determined as previously described using an Oroboros Oxygraph O2k instrument (Oroboros Instruments, Inc., Innsbruck, Austria) ^43–45,76^. All studies utilized permeabilized cells or homogenized tissue in MiR05 buffer containing 10 μM cytochrome c, with or without the sequential addition of TCA cycle substrates to the indicated final concentrations: malate (2 mM), ADP (5 mM), pyruvate (5 mM), glutamate (10 mM), and succinate (10 mM). Final results were adjusted for differences in cell number and mitochondrial mass.

To quantify proton leak, 1.5 x 10^6^ cells in log-phase growth were harvested as described above, placed into Mir05 buffer and permeabilized with digitonin to a final concentration of 10 µg/mL. The following substrates were then sequentially injected to the indicated final concentrations while monitoring OCRs: cytochrome c (10 µM), malate (2 mM), pyruvate (5 mM), glutamate (10 mM), succinate (10 mM), octanoate (0.5 mM), and glycerophosphate (10 mM). OCR (in the absence of ADP) was then designated as “leak” respiration. ADP was then added to a final concentration of 1.25 mM to initiate OXPHOS, followed by the stepwise titration of carbonyl cyanide 3-chlorophenylhydrazone (m-Cl-CCP) (0.25 µM per step) to achieve maximal uncoupled respiration. Leak fraction was calculated as the ratio of leak OCR to maximal OCR.

Rates of ATP synthesis were measured in digitonin-permeabilized cells that were prepared as described above. This was followed by the addition of Magnesium Green (Thermo Fisher) to a final concentration of 1.1 μM and the sequential addition of (palmitate+malate+ADP), glutamate and succinate while also continuously measuring OCRs in parallel ^103,104^. ATP synthesis rates were quantified by measuring changes with the MgG / CaG Filter Set (Oroboros Instruments GmbH, Cat# 44323-01).

### Quantification of pyruvate dehydrogenase activity and fatty acid β oxidation (β-FAO)

Assays for pyruvate dehydrogenase in livers and HBs were performed as previously described ^41^. Briefly, ∼50 mg of snap-frozen tissue was finely minced in one ml of D-MEM lacking pyruvate, glucose and glutamine. 0.5 ml of the suspension was then diluted with equal volume of 2 X assay buffer that also contained 0.15 μCi of [1-^14^C]- labeled pyruvate (7.64 mCi/mmol, PerkinElmer, Waltham, MA). Reactions were performed in sealed tubes with a hanging basket (Kimble-Chase Life Science and Research Products, Rockwood, TN) that contained a glass microfiber filter (Whatman/GE Healthcare, Chicago, IL) soaked in 0.5M KOH. Reactions were incubated for one hr at 37°C and terminated by injecting 0.5 ml of 4M perchloric acid through the rubber cap. The released ^14^CO_2_ was collected for 40 min at 37°C and quantified by scintillation counting of the air-dried filters.

To measure rates of β-FAO, the indicated cell lines were seeded into 24 well plates and grown for 1-2 days to ∼30% confluence. The medium was then removed and one ml of fresh D-MEM+10% FBS was added to each well followed by incubation in either 20% or 1% oxygen for an additional 48 hr. 50 uL of a palmitate solution was added to each well to a final concentration of 100 μM that included 0.5 μCi/well of ^14^C-palmitate (PerkinElmer, Inc. 55 mCi/mmol). This mixture was first solubilized in 10 mg/ml α cyclodextrin prior to adding it to the media. After an additional 2 hr incubation at 37C in their original oxygen concentrations, the medium was removed, placed into Eppendorf tubes and acidified by the addition of concentrated perchloric acid to a final concentration of 0.5 M. The released ^14^C-CO_2_ was captured on 1 cm Whatman paper discs (Sigma-Aldrich) that were soaked in 2M KOH. The disks were then removed and ^14^C-CO_2_ was quantified in a scintillation counter. Results were then normalized to cellular protein content. All assays were performed on 3-4 replicas per condition.

### CV ATPase assays

ETC isolation, non-denaturing gel electrophoresis and *in situ* CV ATPase assays were performed as described previously in detail ^7,43^. Briefly, livers, tumors or *in vitro-*propagated cells were harvested and immediately placed into 7 volumes of ice-cold SET buffer. After cutting tissues into 1-2 mm pieces they or the cells were homogenized and then clarified by centrifugation (600 x g) at 4C for 10 min. After discarding the pellets, the homogenates were again centrifuged (7000 x g) at 4C for 10 min. The pellets containing the mitochondrial fraction were then washed with SET buffer, centrifuged again, snap frozen in liquid nitrogen and stored at −80C. Subsequently, the frozen pellets were re-suspended in 200 μl of HB buffer (300mM-Sucrose, 50mM-potassium phosphate, pH-7.4 plus protease inhibitor cocktail (PMSF, PepA, Leupeptin, Aprotinin). After determining the protein concentrations using a BCA assay (Thermo Fisher), the mitochondria were again pelleted at 7,000 x g for 10 min and then resuspended in 30mM-HEPES, pH 7.4; 150mM potassium acetate and 10% glycerol. An amount of digitonin equal to 8 times that of the total protein weight was then added. Following incubation on ice for 1.5 hr, the complexes were resolved by non-denaturing gel electrophoresis on a 3-12% gradient Native PAGE Novex Bis-Tris gels at 4C (Invitrogen, Inc., Carlsbad, CA). After carefully rinsing the gel 3 times in distilled water and equilibrating it in 20 mM Tris-HCl, pH8.2 for 1.5 hr, *in situ* ATPase activity was determined by adding ∼25 ml of a solution containing 34 mm Tris-glycine, pH 7.8; 14 mM MgSO_4_, 8 mM ATP and 0.2% Pb(NO_3_)_2_, pH 7.8. Imaging was performed after a 24 hr incubation at room temperature in the dark and then daily afterward for up to 5 days. Gel were analyzed with a Bio-Rad ChemiDox^TM^ XRS^+^ imaging system and the bands quantified with the accompanying software package (Bio-Rad, Inc., Hercules, CA)

### Immunoblotting

Tissue lysates were prepared in sodium dodecyl sulfate-lysis buffer as previously described ^43,44^. Immunoblots were developed by using SuperSignal West Pico Chemiluminescent Substrate kit (Thermo Fisher). Antibodies, the vendors from which they were obtained and the dilutions for immunoblotting are listed in Supplementary Table 1).

### Mass Spectrometry

The regions of non-denaturing polyacrylamide gels containing ATPase activity of interest (Figure 2C) were excised for mass spectrometric (MS) analysis by The University of Pittsburgh’s Health Sciences Mass Spectrometry Core as previously described ^76^.. Briefly, gel slices were washed with HPLC water and several changes of 50% acetonitrile (ACN)/25mM ammonium bicarbonate. The bands were then dehydrated with 100% ACN, reduced with 10mM dithiothreitol (DTT) at 56°C for 1 hour, followed by alkylation with 55 mM iodoacetamide (IAA) at room temperature for 45 min in the dark. They were then dehydrated with 100% ACN to remove excess DTT and IAA. Trypsin in 25 mM ammonium bicarbonate was added to a final concentration of 20 ng/μl and incubated overnight at 37°C. The resultant tryptic peptides were extracted with 70% ACN/5% formic acid (FA), vacuum dried, and re-constituted in 20 μl 3%ACN/0.1% formic acid. MS analysis was conducted on a Bruker timsTOF Pro2 instrument coupled to a NanoElute 2 liquid chromatography system. Peptides were loaded onto an Aurora Ultimate C18 column (1.7 µm, 25 cm x 75 µm, IonOpticks) and eluted at 300 nl/min over a 60-minute gradient. The timsTOF Pro2 was set to PASEF scan mode and DDA with a scan range of 100 - 1700 m/z with 10 PASEF ramps. The TIMS was set to a 100 ms ramp and accumulation time (100% duty cycle) with a ramp rate of 9.43 Hz. Linear precursor repetitions were set to a target intensity of 20,000 and a target threshold of 2500 and active exclusion was set to 0.40 min. Collision energy was set to a base of 1.60 1/K0 [V-s/cm2] at 59 eV and 0.60 1/ K0 [V-s/cm2] at 20 eV. An isolation width was set to 2 m/z for <700 m/z and 3 m/z for >800 m/z.

The collected MS data were analyzed using MSFragger V3.8 ^105^ against the mouse protein sequence from SwissProt. The search parameters were set as follows: strict trypsin digestion, nonspecific cleavage, up to missed cleavages 2, carbamidomethylation of cysteine as static modification, oxidization of methionine and protein N-terminal acetylation as variable modification, a maximal mass tolerance of 10 ppm for the precursor ions and 20ppm for the fragment ions, and false detection rate (FDR) was capped at 1%.

e mouse protein sequence from SwissProt. The search parameters were set as follows: strict trypsin digestion, nonspecific cleavage, up to missed cleavages 2, carbamidomethylation of cysteine as static modification, oxidization of methionine and protein N-terminal acetylation as variable modification, a maximal mass tolerance of 10 ppm for the precursor ions and 20ppm for the fragment ions, and false detection rate (FDR) was capped at 1%.

### Statistical analysis of transcriptomic and other data

Transcriptomes (HTSeq-FPKM-UQ files) of relevant cancers and control tissues from The Cancer Genome Atlas (TCGA) were downloaded using the TCGAbiolinks R package. Our own previously published RNAseq data on murine HBs and HCCs were accessed from The National Center for Biotechnology Information (NCBI) Gene Expression database and can be accessed through the Gene Expression Omnibus (GEO) under accession numbers <u>GSE303515</u> and <u>GSE130178</u>, respectively ^43–45^. Survival data were analyzed with GraphPad Prism 7 (GraphPad Software, Inc, San Diego, CA) with different expression level cutoffs being used to define intragroup subsets with high and low levels of expression of particular transcripts. The log-rank (Mantel-Cox) test was used to compile Kaplan-Meier survival curves. ANOVA was applied for multiple comparisons using Fisher least significant difference test. Student 2-tailed *t* test was used for comparing differences between 2 groups.

## Acknowledgements

Computational analyses were supported by the University of Pittsburgh Center for Research Computing and by NIH award number S10OD028483. Additional funding was provided by grants from the NIH (RO1 CA174713), The Rally Foundation (no. 22N42) and a Hyundai Hope on Wheels Scholar grant (all to EVP). Additional support was provided by The UPMC Children’s Hospital Foundation and by a Seed grant from The UPMC Children’s Hospital Research Advisory Committee. Work performed in the Health Sciences Mass Spectrometry Core (RRID:SCR_025222) and services and instruments used in this project were graciously supported, in part, by the University of Pittsburgh and the Office of the Senior Vice Chancellor for Health Sciences.

## Author contributions

HW: designed and performed experiments, assisted with writing the manuscript; JL, CH, ACR, ESG, AT, LH, NM: designed, performed and interpreted experiments; SJM: performed and interpreted MS studies; EVP: designed, interpreted and organized experiments, wrote the paper.

## SUPPLEMENTARY FIGURE LEGENDS

**Supplementary Figure 1.**
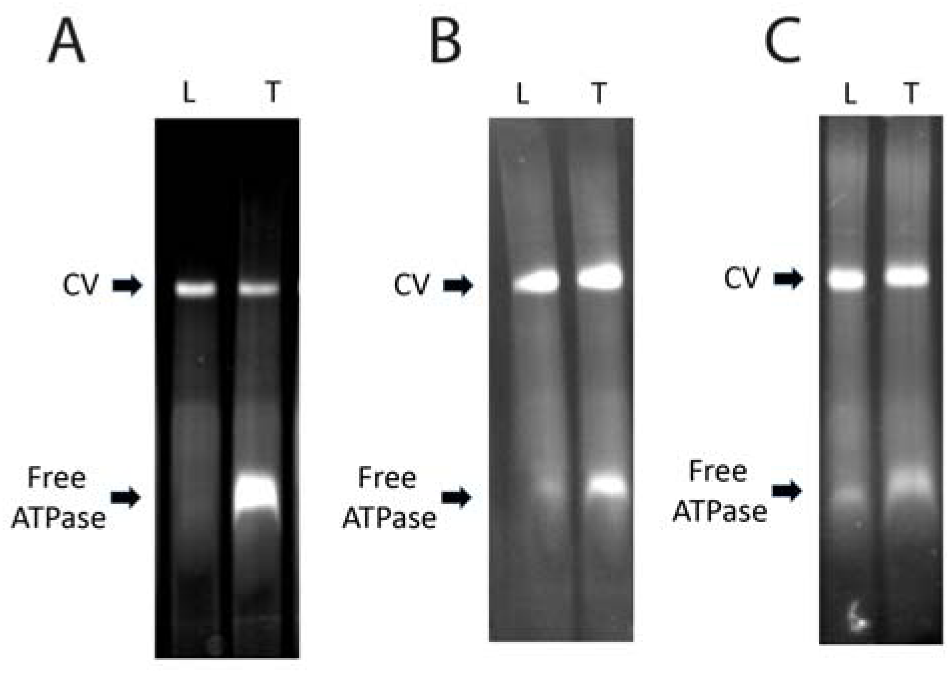
Free ATPase activity remains detectable in BY HBs generated in livers with different genetic backgrounds. (A). Wild-type background (repeat of Figure 1A) (B). *Pdha1-/-* hepatocytes. L: control *Pdha1-/-* livers. T: tumors (C). *Chrebp-/-* hepatocytes. L: control *Chrebp-/-* livers. T: tumors

**Supplementary Figure 2.**
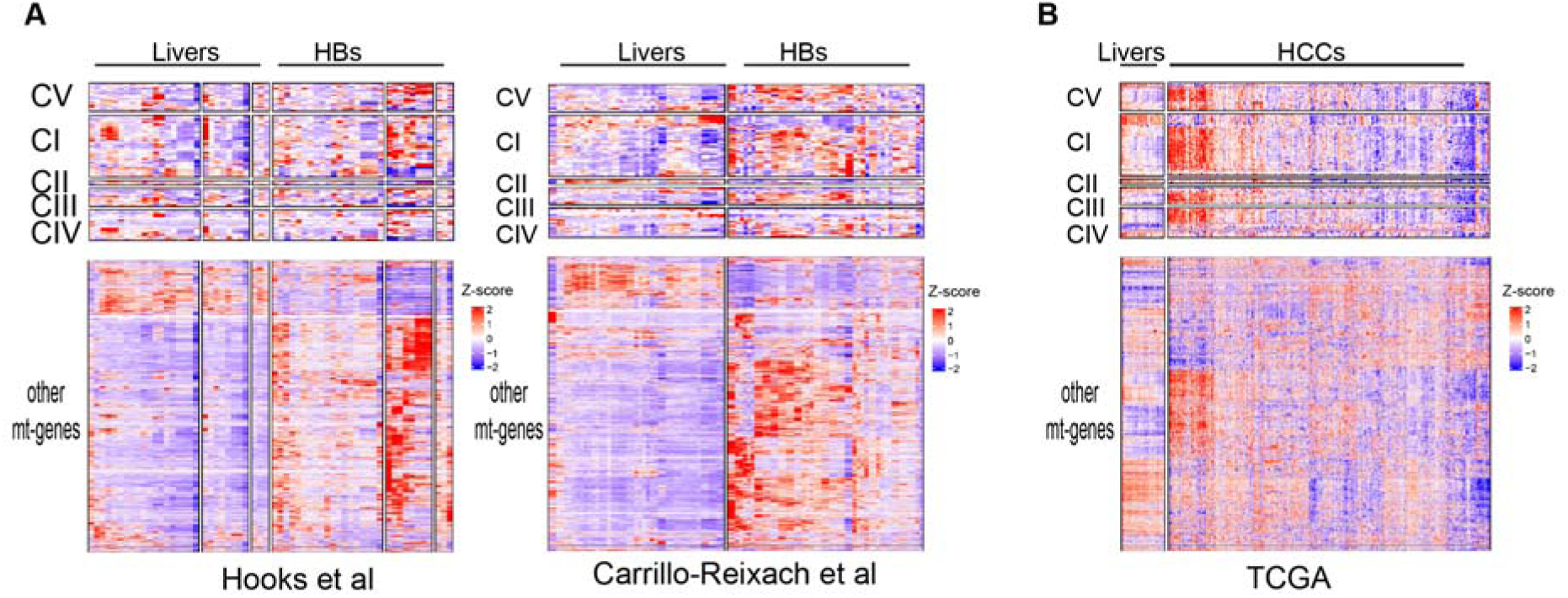
Dysregulation of mitochondrial-specific transcripts in subsets of human HBs and HCCs. (A). HBs. RNAseq data from 2 previously published cohorts of 49 human HBs and matched normal livers were evaluated for their expression of mitochondrial-specific transcripts ^55,56^. The tops of each panel show expression of transcripts encoding subunits for CV and each of the 4 ETC complexes. All other transcripts (∼1100) are shown in the bottom panels. (B). RNAseq studies from human HCCs and matched normal livers from TCGA. Transcripts are arranged as described in (A).

**Supplementary Figure 3.**
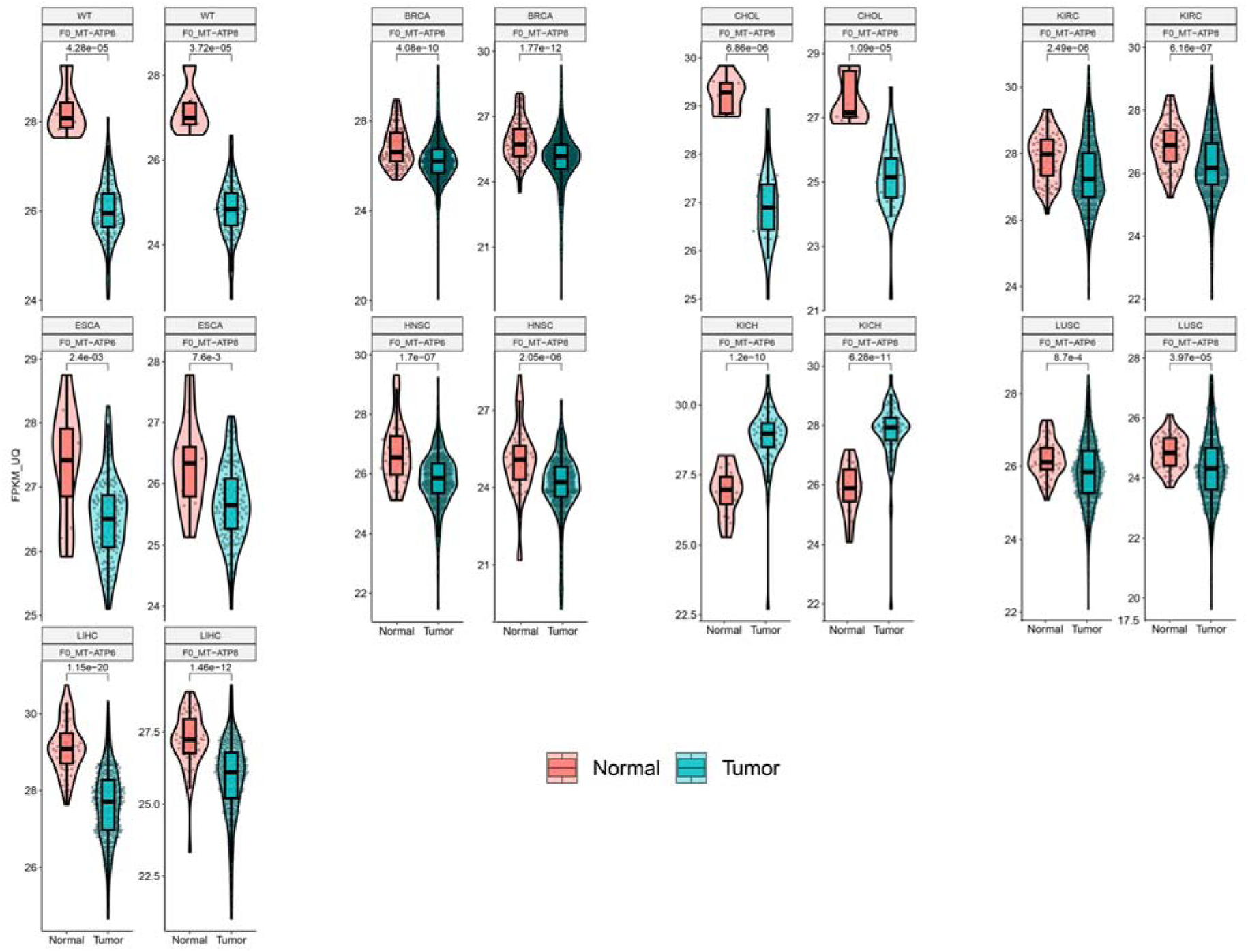
Coordinate down-regulation of *MT-ATP6* and *MT-ATP8* transcripts in select human cancers. RNAseq results from the indicated tumors and matched autologous normal tissues were obtained from the TCGA database and expressed as violin plots. WT: Wilms tumor; BRCA: breast cancer; CHOL: Cholangiocarcinoma; ESCA: esophageal cancer; HNSC: head and neck squamous cell cancer; KIRC: renal clear cell carcinoma; LIHC: hepatocellular carcinoma (HCC); LUSC: lung squamous cell cancer. *: P<0.05, **: P<0.01, ***: P<0.005; ****: P<0.001.

**Supplementary Figure 4.**
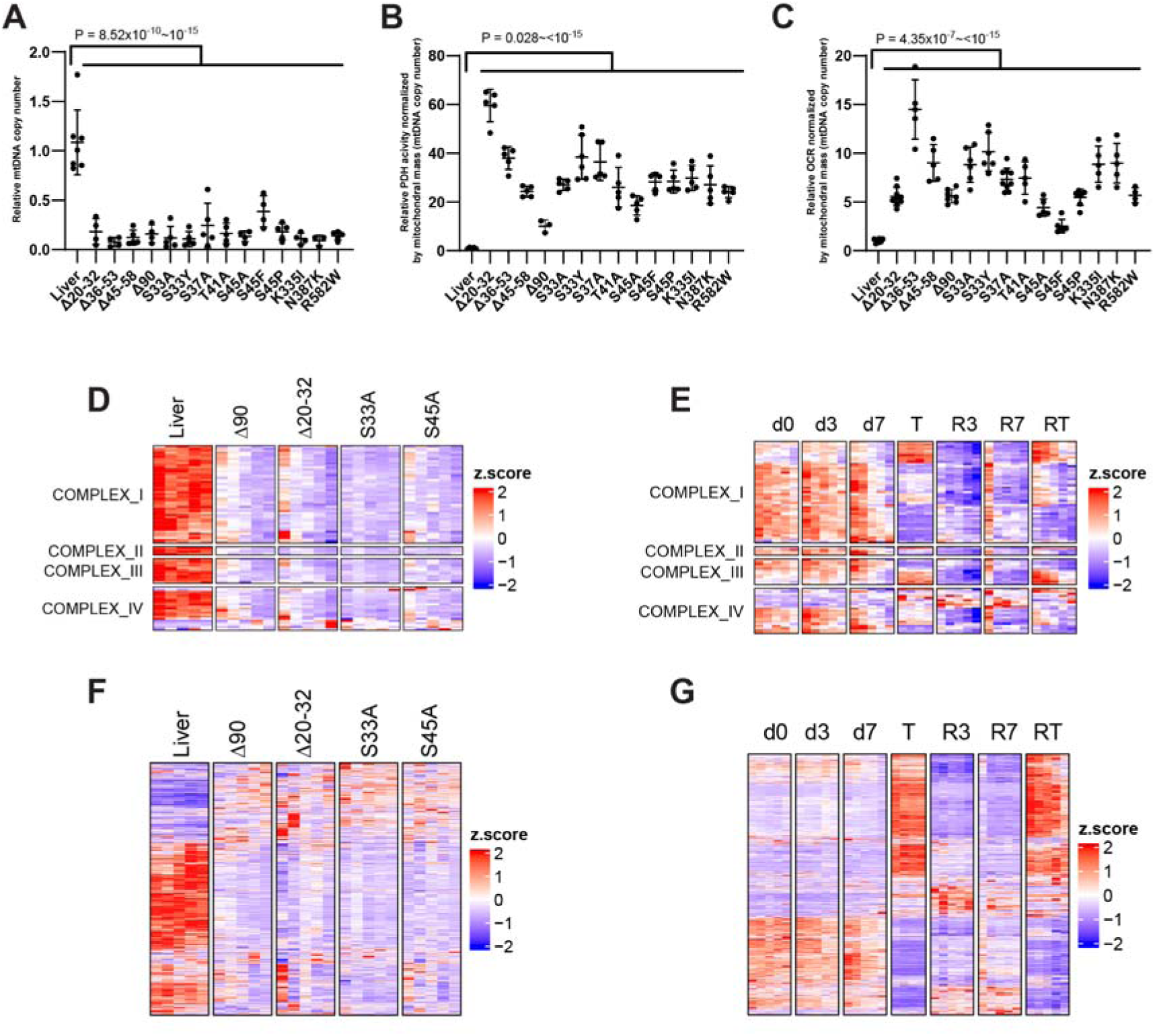
HBs have low mitochondrial mass but high TCA cycle function. (A). mtDNA content of livers and HBs generated by Y^S127A^ + 14 patient-derived β-catenin mutants (including B [Δ90]) ^43^. Each point shows the mean of duplicate TaqMan assays for mtDNA content, performed in triplicate with 2 different sets of mtDNA probes ^43,45^. (B). Complex I activities of the tissues depicted in A measured as OCRs in response to pyruvate, malate, glutamate and ADP. The results were adjusted to account for the mtDNA differences shown in A. OCRs were quantified with an Oroboros respirometer as previously described ^43,45^. (C). Pyruvate dehydrogenase (PDH) activity as measured by the oxidation of 1-^14^C-pyruvate to acetyl-CoA and ^14^CO_2_ ^41,43,44,76^. Results were adjusted to account for mtDNA content differences as in (B). (D). Composite heat map showing relative expression levels of transcripts encoding subunits of CI-CIV of the ETC in livers and 4 sets of HBs from panels A-C. (E). Composite heat map showing relative expression of the transcripts encoding subunits of CI-CIV of the ETC during the course of *MYC*-induced HCC induction, regression and recurrence (Figure 1C). (F). Transcripts encoding mitochondrial proteins (other than those for CI-CV) in a subset of the murine livers and HBs shown in (A-C). (G). Transcripts encoding mitochondrial proteins (other than those for CI-CV) in livers and murine HCCs.

**Supplementary Figure 5.**
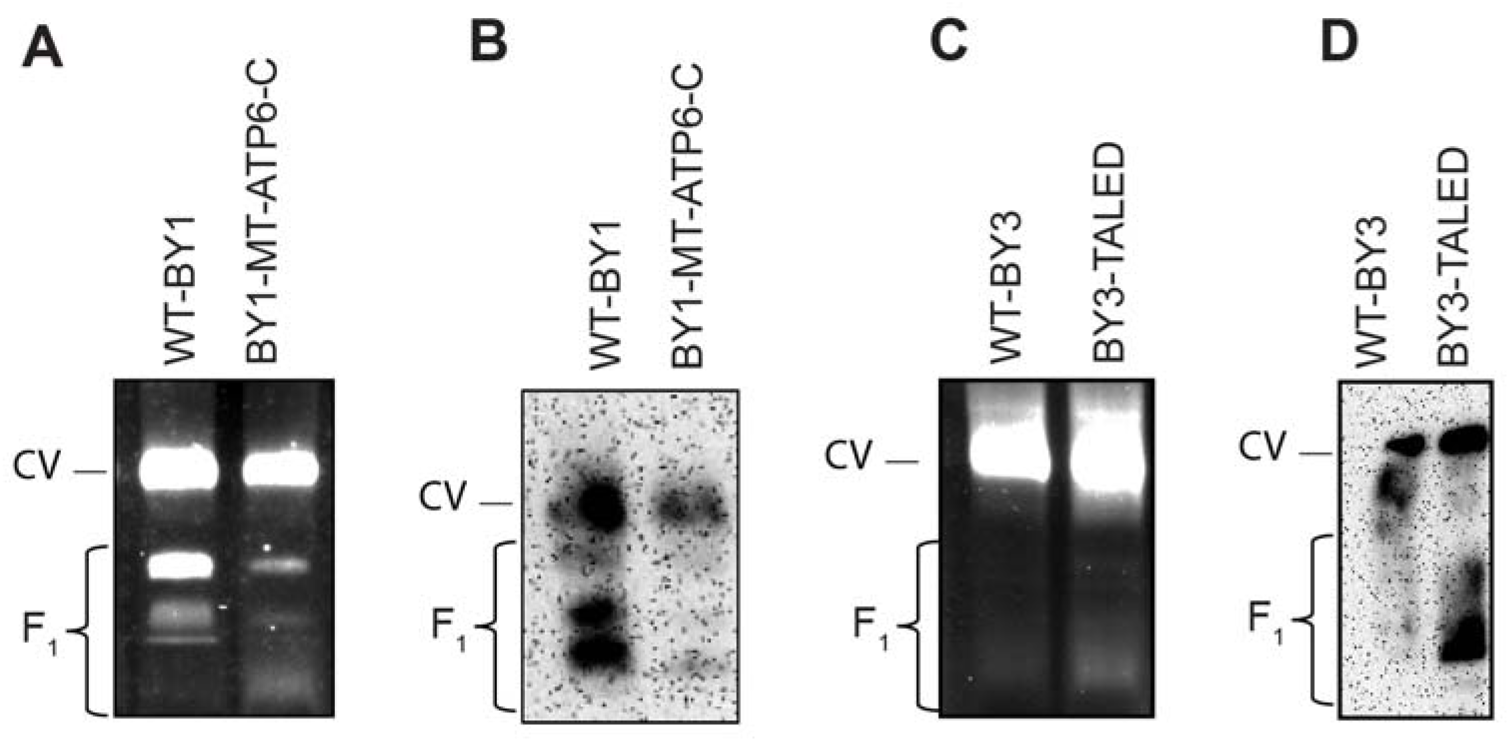
Partial restoration of intact CV by the enforced expression of Mt-atp6-c. (A). *In situ* ATPase assays on BY1 cells showing a ∼80% reduction of free F_1_ following stable expression of the Mt-atp6-c fusion protein (Figure 2H). Numbers below the gel indicate the ratio of free F_1_ ATPase activity relative to total in each lane. (B). A non-denaturing gel identical to that shown in A was transferred to a PVDF membrane and incubated with an anti-Atp5f1a (α subunit) antibody. (C). NDGE and *in situ* ATPase assays showing that TALED-mediated targeting of BY3 cells increased the amount of free F_1_. (D). A non-denaturing gel identical to that shown in C was immuno-blotted with an anti-α subunit antibody as described in (B).

**Supplementary Figure 6.**
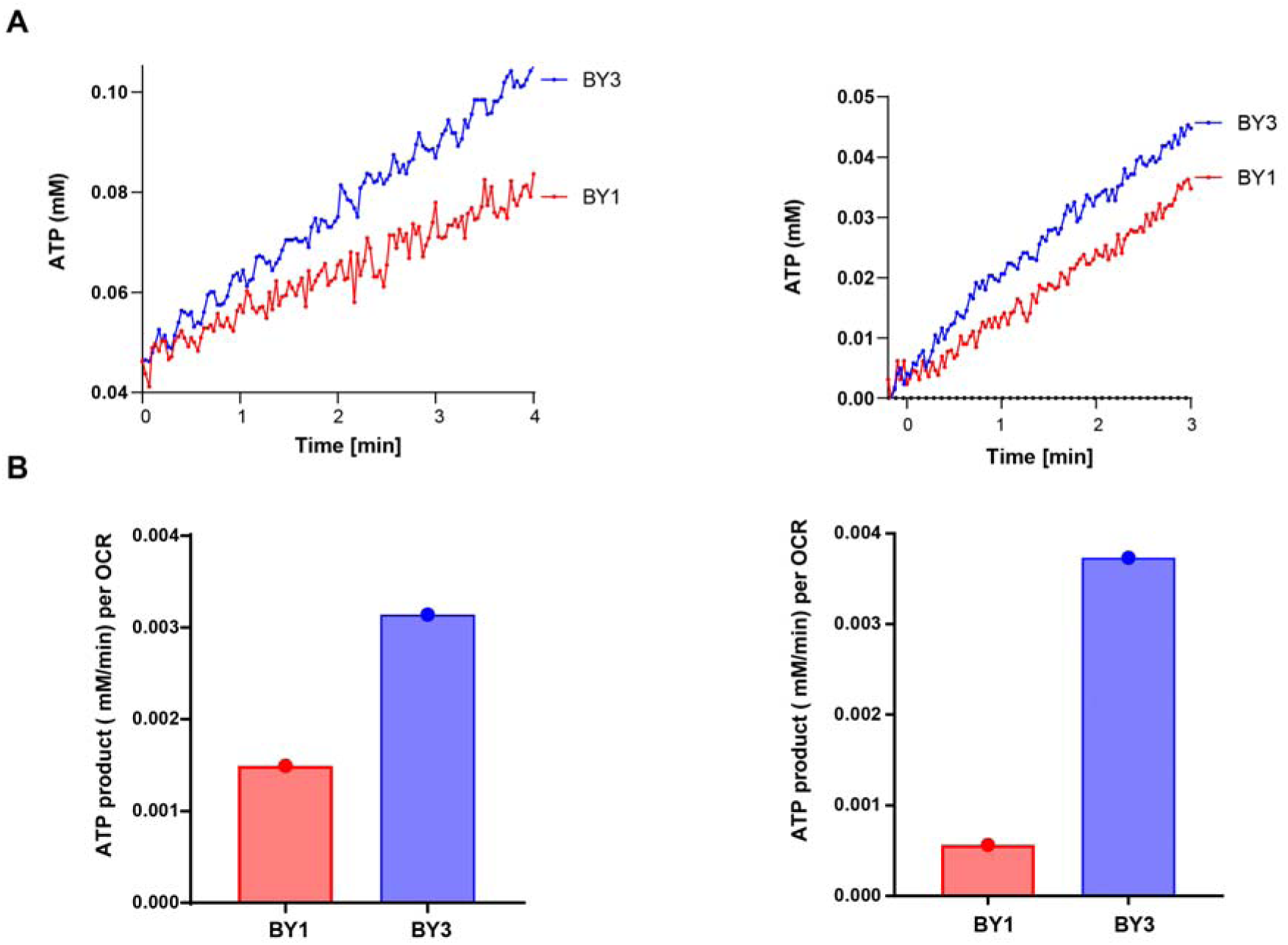
Inefficient ATP generation by BY1 cells. (A). Rates of ATP accumulation are greater in BY3 cells. Equal numbers of BY1 and BY3 cells were permeabilized with digitonin. ATP accumulation rates in response to the addition of pyruvate, malate, glutamate and succinate was measured in real time using a Mg++ Green assay ^103,104^. Shown are the results of 3 separate experiments performed at different times. (B). ATP production from (A) normalized to OCRs measured in parallel response to the addition of pyruvate, malate, glutamate and succinate and ADP from each of the experiments shown in (A).

## SUPPLEMENTARY TABLES

**Supplementary Table 1.** Antibodies, vendors and dilutions used.

| Name of Antibody | Vendor | Catalog No. | Dilution used |
| --- | --- | --- | --- |
| Anti-Mt-Atp6 | Cell Signaling Technology, Inc | 70262 | 1:1000 |
| Anti-Atp5fa1 | Cell Signaling Technology, Inc | 18023 | 1:1000 |
| Anti-V5 epitope | ThermoFisher | 46-07-05 | 1:5000 |
| Anti-Pdha1 | Cell Signaling Technology, Inc | 3205 | 1:1000 |
| Anti-Gapdh | Sigma | G8795 | 1:10000 |
| Anti-β-action | Cell Signaling Technology, Inc | 3700 | 1:1000 |
| Anti mouse HRP | Cell Signaling Technology, Inc | 7076 | 1:10000 |
| Anti rabbit HRP | Cell Signaling Technology, Inc | 7074 | 1:5000 |

## SUPPLEMENTARY FILES

**Supplementary File 1 Incidence of TALED-induced mutations generated in BY3 cells.**

